# The Chromatin Landscape of Pathogenic Transcriptional Cell States in Rheumatoid Arthritis

**DOI:** 10.1101/2023.04.07.536026

**Authors:** Kathryn Weinand, Saori Sakaue, Aparna Nathan, Anna Helena Jonsson, Fan Zhang, Gerald F. M. Watts, Zhu Zhu, Accelerating Medicines Partnership Program: 5 Rheumatoid Arthritis and Systemic Lupus Erythematosus (AMP RA/SLE) Network, Deepak A. Rao, Jennifer H. Anolik, Michael B. Brenner, Laura T. Donlin, Kevin Wei, Soumya Raychaudhuri

## Abstract

Synovial tissue inflammation is the hallmark of rheumatoid arthritis (RA). Recent work has identified prominent pathogenic cell states in inflamed RA synovial tissue, such as T peripheral helper cells; however, the epigenetic regulation of these states has yet to be defined. We measured genome-wide open chromatin at single cell resolution from 30 synovial tissue samples, including 12 samples with transcriptional data in multimodal experiments. We identified 24 chromatin classes and predicted their associated transcription factors, including a *CD8*+ *GZMK*+ class associated with EOMES and a lining fibroblast class associated with AP-1. By integrating an RA tissue transcriptional atlas, we found that the chromatin classes represented ‘superstates’ corresponding to multiple transcriptional cell states. Finally, we demonstrated the utility of this RA tissue chromatin atlas through the associations between disease phenotypes and chromatin class abundance as well as the nomination of classes mediating the effects of putatively causal RA genetic variants.

## Introduction

Rheumatoid arthritis (RA) is a chronic autoimmune disease that affects roughly one percent of the population^1^. In RA, the synovial joint tissue is infiltrated by immune cells that interact with stromal cells to sustain a cycle of inflammation. Untreated, RA can lead to joint destruction, disability, and a reduction in life expectancy^2^. The heterogeneous clinical features of RA, including differences in cyclic citrullinated peptide antibody autoreactivity^3^, underlying genetics^4, 5^, and response to targeted therapies^6–10^, render it challenging to construct generic treatment plans that will be effective for most patients.

Recent studies have taken advantage of single cell technologies to define key cell populations that are present and expanded in RA tissue inflammation^11–14^, demonstrating both the heterogeneous nature of tissue inflammation and the promise to identify novel targeted therapeutics for RA. Our recent AMP-RA reference study^12^ comprehensively classified pathogenic transcriptional cell states within synovial joint tissue using single cell CITE-seq technology^15^, which simultaneously measures mRNA and surface protein marker expression in a single cell. Within 6 broad cell types (B/plasma, T, NK, myeloid, stromal [fibroblasts/mural], and endothelial), the study defined 77 fine-grain cell states. Many of these cell states have been previously shown to be associated with RA pathology: for example, CD4+ T peripheral helper cells (TPH)^11, 13^, HLA-DR^hi^ sublining fibroblasts^11^, proinflammatory IL1B+ monocytes^11^, and age-associated B cells (ABC)^11, 16^. However, we have a limited understanding of the chromatin accessibility profiles that underlie these pathogenic synovial tissue cell states.

Open chromatin at critical *cis*-regulatory regions allows essential transcription factors (TFs) to access DNA and epigenetically regulate gene expression^17^. Chromatin accessibility is a necessary, but not sufficient, condition for RNA polymerases to produce transcripts at gene promoters^18^. Therefore, one possibility is that each transcriptional cell state has its own unique chromatin profile^19^, which we will denote as a chromatin class. Alternatively, multiple transcriptional cell states could share a chromatin class if the cell states were dynamically transitioning from one to another in response to external stimuli without altering the chromatin landscape^19^. In RA, those external stimuli could be cytokines that activate TFs to induce expression of key genes and drive pathogenic cell states^20^. For example, NOTCH3 signaling propels transcriptional programs coordinating the transformation from perivascular fibroblasts to inflammatory sublining fibroblasts^21^. Similarly, exposure to TNF and interferon gamma transforms monocytes into inflammatory myeloid cells^22^.

Here, we characterized synovial cells with unimodal single cell ATAC-seq (scATAC) and multimodal single nuclear ATAC-seq (snATAC) and RNA-seq (snRNA) technologies to compare chromatin classes to transcriptional cell states (**Fig. 1a**). These results support a model of open chromatin superstates shared by multiple fine-grain transcriptional cell states. We show these superstates may be regulated by key TFs and associated with clinical and genetic factors in the pathology of RA (**Fig. 1a**).

**Fig. 1.**
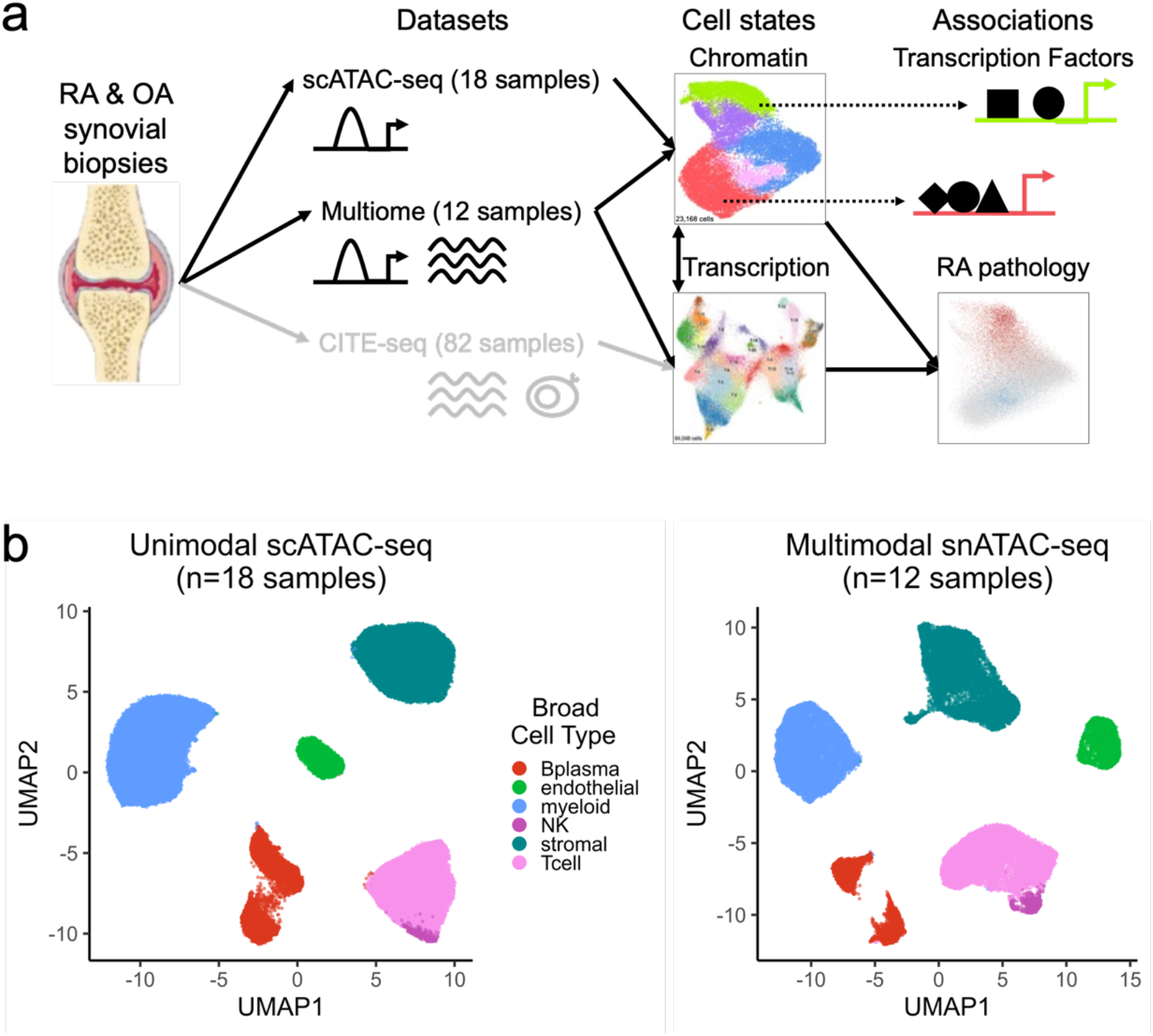
Study overview and open chromatin broad cell type identification. **a.** Study overview. Synovial biopsies from RA and OA patients were utilized for unimodal scATAC-seq, multimodal snATAC-seq + snRNA-seq experiments. CITE-seq was performed in the AMP-RA reference study^12^. We defined chromatin classes using the unimodal and multimodal ATAC data and compared them with AMP-RA transcriptional cell states^12^ classified onto the multiome cells. We further defined transcription factors likely regulating these chromatin classes and found putative links to RA pathology by associating the classes to RA clinical metrics, RA subtypes, and putative RA risk variants. **b.** Open chromatin broad cell type identification in unimodal scATAC-seq datasets (**left**) and multimodal snATAC-seq datasets (**right**), processed separately.

## Results

### Unimodal scATAC and multimodal snATAC synovial tissue datasets

We obtained synovial biopsies from 25 people with RA and 5 with osteoarthritis (OA) and disaggregated cells using well-established protocols from the AMP-RA/SLE consortium^23^ (**Methods**). We conducted unimodal scATAC-seq on samples from 14 RA patients and 4 OA patients and multimodal snATAC-/snRNA-seq on samples from 11 RA patients and 1 OA patient (**Supplementary Table 1**). Applying stringent ATAC quality control, we retained cells with >10,000 reads, >50% of those reads falling in peak neighborhoods, >10% of reads in promoter regions, <10% of reads in the mitochondrial chromosome, and <10% of reads falling in the ENCODE blacklisted regions^24^ (**Methods**; **Supplementary Fig. 1a-b**). We further required that cells from the multimodal data passed stringent quality control for both the snRNA and snATAC (**Supplementary Fig. 1c**). After additional QC within individual cell types combining both technologies, the final dataset contained 86,994 cells from 30 samples (median: 2,990 cells/sample) (**Supplementary Fig. 1d-e**). For consistency, we called a set of 132,520 consensus peaks from unimodal scATAC data to be used for all analyses (**Methods**). We observed that 95% of the called peaks overlapped ENCODE candidate *cis*-regulatory elements (cCREs)^25^ and 17% overlapped promoters^26^, suggesting highly accurate peak calls (**Supplementary Fig. 1f**).

### Defining RA broad cell types by clustering ATAC datasets

To assign each ATAC cell to a broad cell type, we clustered the unimodal scATAC and multimodal snATAC datasets independently (**Methods**). In both instances, we defined six cell types that we annotated based on the chromatin accessibility of “marker peaks,” or peaks in cell type marker gene promoters (**Methods**; **Fig. 1b**). We identified T cells (*CD3D* and *CD3G*), NK cells (*NCAM1* and *NCR1*), B/plasma cells (*MS4A1* and *TNFRSF17*), myeloid cells (*CD163* and *C1QA*), stromal cells (*PDPN* and *PDGFRB*), and vascular endothelial cells (*VWF* and *ERG*) (**Supplementary Fig. 1g-j**). In the multimodal data, we observed consistent peak accessibility and RNA expression for marker genes in these cell types (**Supplementary Fig. 1k-m**).

We combined ATAC cells from multimodal and unimodal technologies and then created datasets for each of the broad cell types. For cell types with more than 1,500 cells, we applied Louvain clustering to a shared nearest neighbor graph based on batch-corrected^27^ principal components of chromatin accessibility to define fine-grain chromatin classes (**Methods**).

### RA T cell chromatin classes

We first examined the accessible chromatin for 23,168 T cells across unimodal and multimodal ATAC datasets. Louvain clustering defined 5 T cell chromatin classes, denoted as T_A_ for T cell ATAC, across 30 samples (**Fig. 2a**; **Supplementary Fig. 2a**). In the T_A_-2: CD4+ PD-1+ TFH/TPH chromatin class, we observed high promoter accessibility and gene expression for *PD-1* (*PDCD1*) and *CTLA4*, known marker genes for T follicular helper (TFH)/T peripheral helper (TPH) cells (**Fig. 2b**; **Supplementary Fig. 2b**). A known expanded pathogenic cell state in RA, TFH/TPH cells help B cells respond to inflammation^11, 13^. The T_A_-3: CD4+ IKZF2+ Treg cluster had high accessibility and expression for *IKZF2* (Helios), which is known to stabilize the inhibitory activity of regulatory T cells^28^ (Tregs) (**Fig. 2b**). We also observed open chromatin regions at both the *FOXP3* transcription start site (TSS) as well as the downstream Treg-specific demethylated region^29^ (TSDR) specifically for T_A_-3 (**Supplementary Fig. 2c**); *FOXP3* was also expressed exclusively in T_A_-3 cells (**Supplementary Fig. 2b**). We found one more predominantly CD4+ T cell class, T_A_-1: CD4+ IL7R+, with high expression and accessibility for *IL7R*, encoding the CD127 protein. This marker is typically lost with activation, suggesting that T_A_-1 is a population of unactivated naive or memory T cells, as further evidenced by *SELL* and *CCR7* expression (**Fig. 2B**; **Supplementary Fig. 2b**). The T_A_-0: CD8+ GZMK+ cluster was marked by *GZMK* and *CRTAM* peak accessibility and gene expression (**Fig. 2b**; **Supplementary Fig. 2b**); a similar population has been shown to be expanded in RA and a major producer of inflammatory cytokines^11, 30^. We found another primarily CD8+ group of T cells, the T_A_-4: CD8+ PRF1+ cytotoxic cluster, which had high accessibility for the *PRF1* promoter and expression for the *PRF1*, *GNLY*, and *GZMB* genes (**Fig. 2b**; **Supplementary Fig. 2b**).

**Fig. 2.**
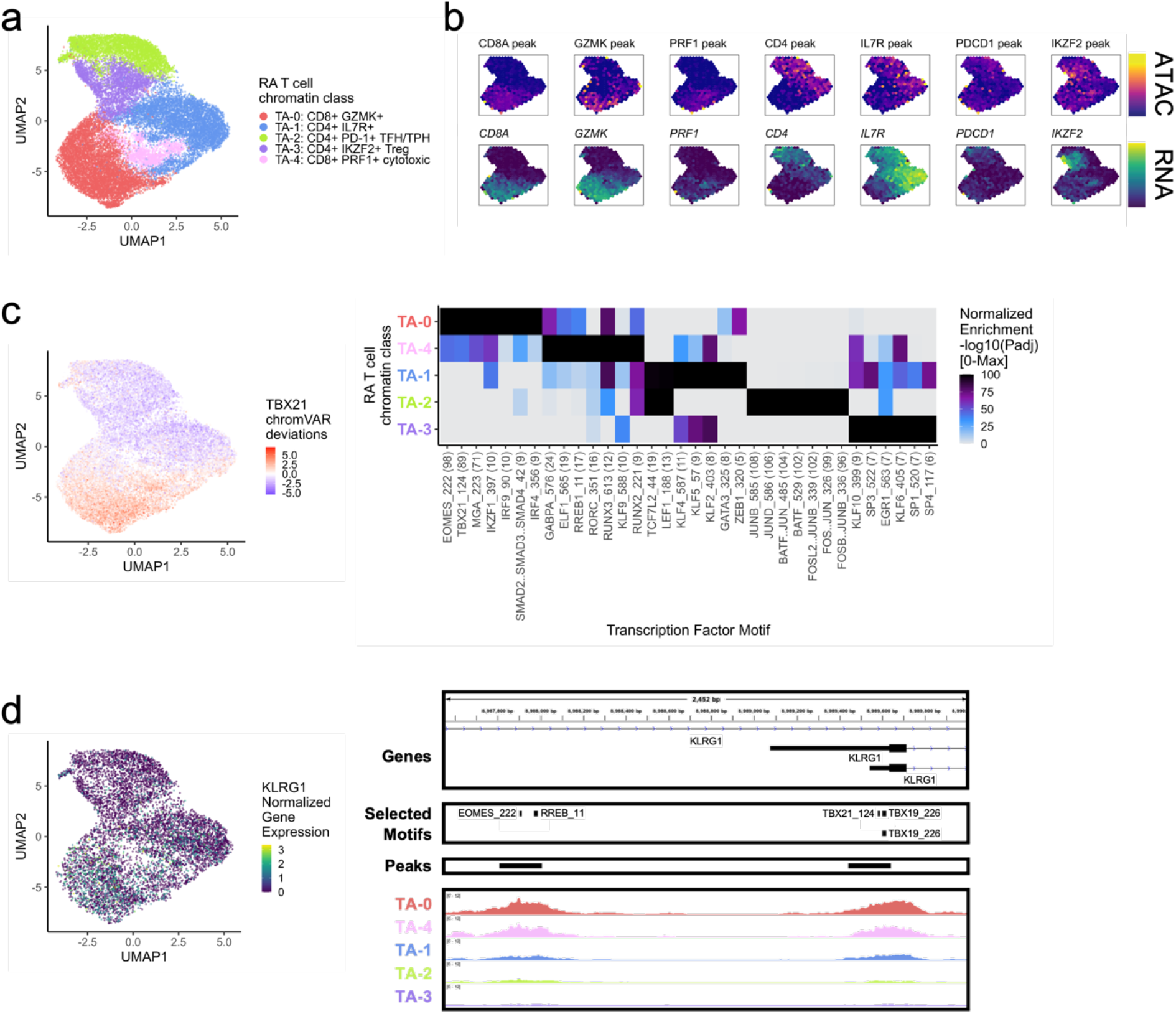
RA T cell chromatin classes. **a.** UMAP colored by 5 T cell chromatin classes defined from unimodal scATAC and multimodal snATAC cells. **b.** Binned normalized marker peak accessibility (**top**) and gene expression (**bottom**) for multiome snATAC cells on UMAP. **c.** UMAP colored by chromVAR^31^ deviations for the TBX21 motif (**left**). Most significantly enriched motifs in marker peaks per T cell chromatin class (**right**). To be included per class, motifs had to be enriched in the class above a minimal threshold and corresponding TFs had to have at least minimal expression in snRNA (**Methods**). Color scale normalized per motif across classes with max -log10(p_adj_) value shown in parentheses in motif label. P-values were calculated via hypergeometric test in ArchR^32^. **d.** UMAP colored by *KLRG1* normalized gene expression in multiome cells (**left**). *KLRG1* locus (chr12:8,987,550-8,990,000) with selected isoforms, motifs, open chromatin peaks, and chromatin accessibility reads from unimodal and multimodal ATAC cells aggregated by chromatin class and scaled by read counts per class (**Methods**) (**right**).

Since T cells are primarily defined by CD4 and CD8 lineages that are not thought to cross-differentiate^33^, we next examined whether the chromatin classes strictly segregated by CD4 or CD8 promoter peak accessibility. We observed that each chromatin class, while largely showing accessibility for only one lineage’s promoter, also includes some cells with accessibility for the other lineage’s promoter (**Supplementary Table 2**). For example, cytotoxic T cells in T_A_-4 were more likely to have an accessible CD8A promoter, but also included a minority of cells with accessibility at the CD4 promoter. Therefore, we assessed which promoter peaks were associated with a specific lineage. While accounting for chromatin class, donor, and read depth, we ran a logistic regression model over all cells relating each promoter peak’s openness to CD4/CD8A promoter peak accessibility status: 1 for open CD4 and closed CD8A, -1 for open CD8A and closed CD4, or 0 otherwise (**Methods**). We only found 93 out of 16,383 promoter peaks significantly associated to a lineage’s promoter accessibility, with 29 associating to CD4 and 64 to CD8A, at FDR<0.20 (**Supplementary Table 3**). This suggested that lineage is important for a small subset of genes’ local promoter chromatin environment, such as *IL6ST* in CD4 T cells and *CRTAM* in CD8 T cells, and for those lineage-specific loci, they segregate by chromatin class as expected (**Methods**; **Supplementary Figure 2d**). However, the majority of promoters appeared to be more specifically accessible within their chromatin classes across lineages. This might suggest that the corresponding gene’s function was critical for the class definition, as highlighted by functional genes such as *PRF1* that is expressed in both cytotoxic CD4 and CD8 T cells^34^ as well as the homing gene *CCR7* that acts across both lineages^35^.

We next determined TFs potentially regulating these T cell chromatin classes by calculating TF motif enrichments^31^ per class marker peaks^32^ whose TFs are at least minimally expressed within that class (**Methods**). In the primarily CD8+ classes, T_A_-0: CD8+ GZMK+ and T_A_-4: CD8+ PRF1+ cytotoxic, we found EOMES (p_adj_=7.44e-99, 8.12e-44, respectively) and T-bet (TBX21) (p_adj_=4.92e-90, 2.75e-38, respectively) motifs preferentially enriched (**Fig. 2c**); the corresponding TFs are known to drive memory and effector CD8+ cell states^36^. Furthermore, we found both motifs in the promoter of *KLRG1*, a gene found in CD8+ effector T cells that might participate in the effector-to-memory transition^37^ (**Fig. 2d**). The cytotoxic T_A_-4 class was also enriched for RUNX3^38^ motifs (p_adj_=5.81e-13) (**Fig. 2c**). Within the T_A_-2: CD4+ PD-1+ TFH/TPH class, we observed high enrichments for AP-1 motifs, especially BATF (p_adj_=3.31e-103), which promotes expression of key programs in TFH cells^39^ (**Fig. 2d**). We found TCF7 and LEF1 motifs^40^ within the unactivated T_A_-1: CD4+ IL7R+ cluster (p_adj_=1.14e-10, 3.97e-13, respectively; **Fig. 2d**).

### RA stromal chromatin classes

Next, we analyzed 24,307 stromal cells (**Methods**). With Louvain clustering, we partitioned the cells into 4 open chromatin classes: lining fibroblasts (S_A_-1) along the synovial membrane, sublining fibroblasts (S_A_-0, S_A_-2) filling the interstitial space, and mural cells (S_A_-3) adjacent to blood vessels^41^ (**Fig. 3a**; **Supplementary Fig. 3a**). The most abundant sublining cluster, S_A_-0: CXCL12+ HLA-DR^hi^ sublining fibroblasts, was a proinflammatory cluster marked by *CXCL12*, *HLA-DRA*, and *CD74* accessibility and expression; S_A_-0 also expressed *IL6,* which is an established RA drug target^7, 8^ (**Fig. 3b**; **Supplementary Fig. 3b**). The S_A_-2: CD34+ MFAP5+ sublining fibroblast class had accessible promoter peaks, where available, for the expressed *CD34*, *MFAP5*, *PI16*, and *DPP4* genes, previously reported to represent a progenitor-like fibroblast state shared across tissue types^42–44^ (**Fig. 3b**; **Supplementary Fig. 3b**). The S_A_-1: PRG4+ lining fibroblast chromatin class was characterized with high accessibility and expression of *PRG4* and *CRTAC1* (**Fig. 3b**; **Supplementary Fig. 3b**). We also observed high expression of *MMP1* and *MMP3*, matrix metalloproteinases responsible for extracellular matrix (ECM) destruction^45^, within S_A_-1 (**Supplementary Fig. 3b**). Finally, we found a mural cell cluster, S_A_-3: MCAM+ mural, with both gene expression and promoter peak accessibility for *MCAM* and *NOTCH3* (**Fig. 3b**; **Supplementary Fig. 3b**). In RA, NOTCH3 signaling from the endothelium acts primarily on mural cells, which in turn stimulate sublining fibroblasts along a spatial axis^21^ as seen in the decreasing NOTCH3 gene expression from S_A_-3, S_A_-0, S_A_-2, to S_A_-1 in the multiome cells (**Supplementary Fig. 3b**). Knockout of *NOTCH3* has been shown to reduce inflammation and joint destruction in mouse models^21^.

**Fig. 3.**
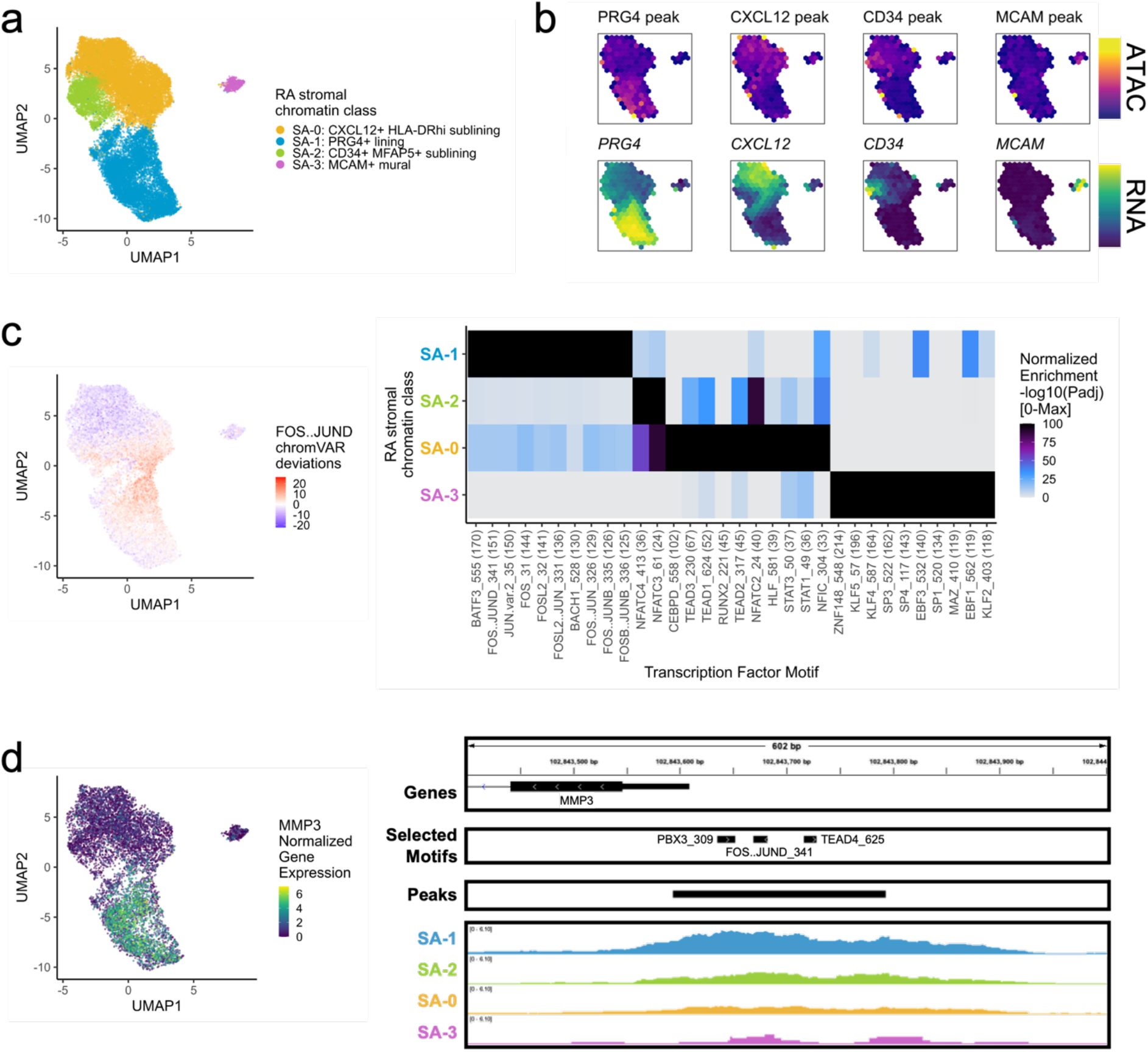
RA stromal chromatin classes. **a.** UMAP colored by 4 stromal chromatin classes defined from unimodal scATAC and multimodal snATAC cells. **b.** Binned normalized marker peak accessibility (**top**) and gene expression (**bottom**) for multiome snATAC cells on UMAP. **c.** UMAP colored by chromVAR^31^ deviations for the FOS..JUND motif (**left**). Most significantly enriched motifs in marker peaks per stromal chromatin class (**right**). To be included per class, motifs had to be enriched in the class above a minimal threshold and corresponding TFs had to have at least minimal expression in snRNA (**Methods**). Color scale normalized per motif across classes with max -log10(p_adj_) value shown in parentheses in motif label. P-values were calculated via hypergeometric test in ArchR^32^. **d.** UMAP colored by *MMP3* normalized gene expression (**left**). *MMP3* locus (chr11:102,843,400-102,844,000) with selected isoforms, motifs, open chromatin peaks, and chromatin accessibility reads from unimodal and multimodal ATAC cells aggregated by chromatin class and scaled by read counts per class (**Methods**) (**right**).

DNA methylation and chromatin accessibility work in tandem to define cell-type-specific gene regulation through silencing CpG-dense promoters and repressing methylation-sensitive TF binding^46^. Methylation changes have been previously described between cultured fibroblast cell lines from RA and OA patients^47, 48^. Thus, we wondered if a specific subset of fibroblasts might be the source of these differentially methylated regions (DMRs). Using a published set of DMRs for RA versus OA synovial fibroblast cell lines^47^, we defined a per-cell score of peak accessibility associated to hypermethylated (positive) or hypomethylated (negative) loci in RA (**Methods**). The sublining fibroblasts in S_A_-0 were enriched for hypomethylated regions (Wilcoxon S_A_-0 cells versus rest one-sided p=0), suggesting that the RA synovial fibroblast DMRs were relatively enriched for putatively functional chromatin accessible regions specifically in sublining fibroblasts (**Supplementary Fig. 3c**). These results proposed the possibility of epigenetic memory retention even after multiple cell line passages^49^, as sublining fibroblasts, particularly HLA-DR^hi^ and CD34^-^ fibroblasts, are expanded in RA relative to OA in synovial tissue samples^11^.

Next, we investigated which TFs were putatively driving these chromatin classes (**Fig. 3c**). AP-1 motifs such as FOS::JUND were most significantly enriched in the S_A_-1 lining class (p_adj_=9.29e-152; **Fig. 3c**). These TFs are known to play many roles in RA and specifically regulate *MMP1* and *MMP3* promoters^49, 50^ (**Fig. 3d**). The progenitor-like sublining S_A_-2 class harbored NFATC motifs, such as NFATC4 (p_adj_=2.89e-36; **Fig. 3c**). In the S_A_-0: CXCL12+ HLA-DR^hi^ sublining chromatin class, we found TEAD1^51^ (p_adj_=2.86e-52; **Fig. 3c**) and STAT1/3 TF motif enrichments (p_adj_=3.34e-37, 4.27e-38, respectively; **Fig. 3c**), the later likely regulating the JAK/STAT pathway responsible for proinflammatory cytokine activation central to RA clinical activity^9, 52^. Finally, S_A_-3: MCAM+ mural cells were enriched for KLF2^53, 54^ and EBF1^55, 56^ motifs (p_adj_=4.94e-119, 1.83e-119, respectively; **Fig. 3c**).

### RA myeloid chromatin classes

We classified 25,691 myeloid cells into 5 chromatin classes (**Fig. 4a**; **Supplementary Fig. 4a**). The first cluster, M_A_-2: LYVE1+ TIMD4+ TRM, is a tissue-resident macrophage (TRM) cluster that had RNA and ATAC signal at *LYVE1*, a perivascular localization marker^14^, and *TIMD4*, a scavenger receptor^14^ (**Fig. 4b**; **Supplementary Fig. 4b**). We found another TRM cluster, M_A_-0: F13A1+ MARCKS+ TRM, with high accessibility and expression at *F13A1* and *MARCKS*, both known to be expressed in macrophages^57, 58^ (**Fig. 4b**; **Supplementary Fig. 4b**). The M_A_-1: FCN1+ SAMSN1+ infiltrating monocytes had accessibility and expression for *FCN1*, *PLAUR*, *CCR2*, and *IL1B*, similar to an expanded proinflammatory population in a previous RA study^11^ (**Fig. 4b**; **Supplementary Fig. 4b**). The M_A_-4: SPP1+ FABP5+ intermediate class likely arose from bone-marrow-derived macrophages^59^ with its high accessibility and expression for *SPP1* (**Fig. 4b**); bone-marrow-derived macrophages are known be abundant in active RA and induce proinflammatory cytokines/chemokines^14, 60^. Finally, we found the M_A_-3: CD1C+ AFF3+ DC chromatin class with expression markers *CD1C*, *AFF3*, *CLEC10A*, and *FCER1A*, whose corresponding promoter peaks generally showed more promiscuity across classes (**Fig. 4b**; **Supplementary Fig. 4b**).

**Fig. 4.**
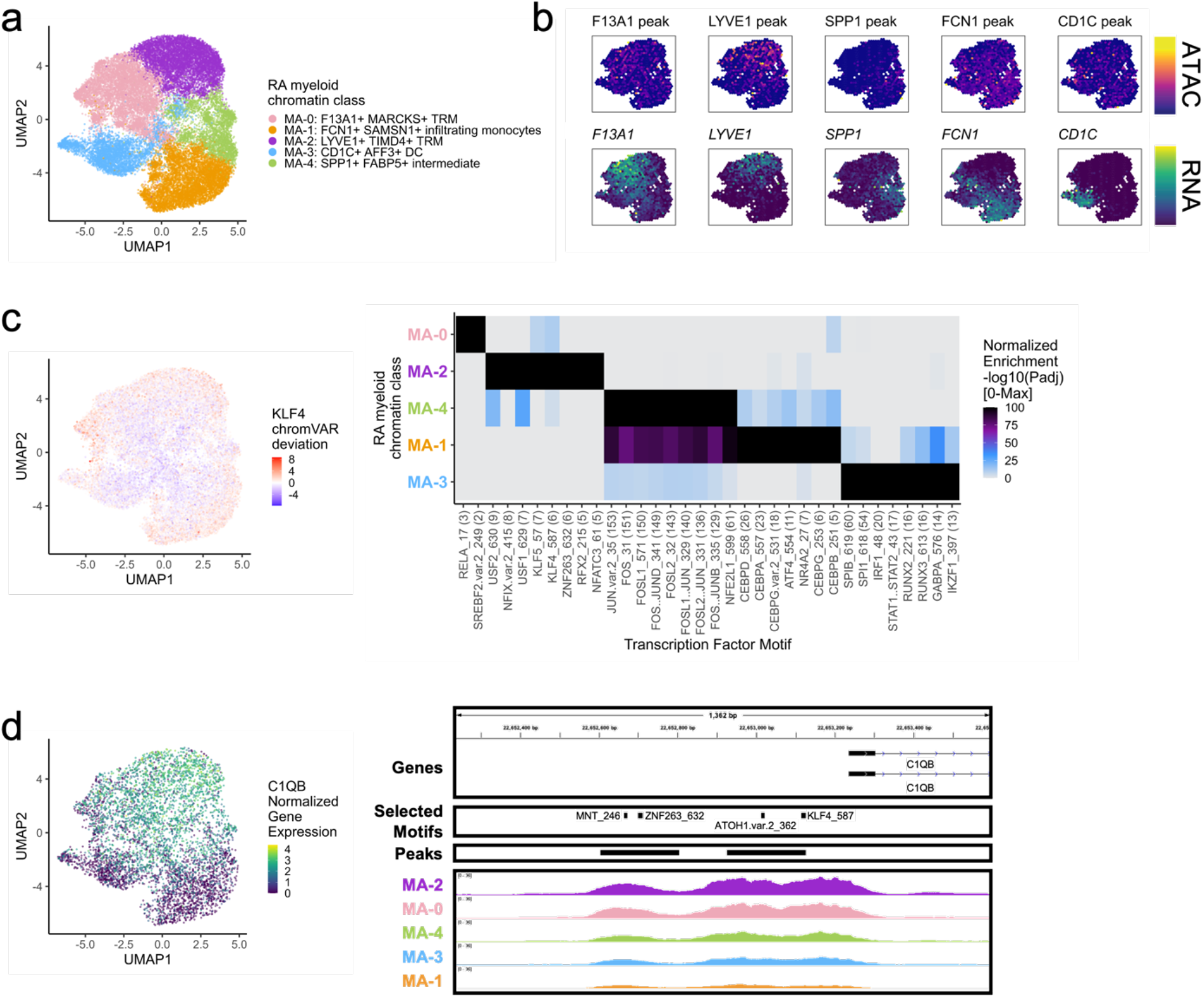
RA myeloid chromatin classes. **a.** UMAP colored by 5 myeloid chromatin classes defined from unimodal scATAC and multimodal snATAC cells. **b.** Binned normalized marker peak accessibility (**top**) and gene expression (**bottom**) for multiome snATAC cells on UMAP. **c.** UMAP colored by chromVAR^31^ deviations for the KLF4 motif (**left**). Most significantly enriched motifs in marker peaks per myeloid chromatin class (**right**). To be included per class, motifs had to be enriched in the class above a minimal threshold and corresponding TFs had to have at least minimal expression in snRNA (**Methods**). Color scale normalized per motif across classes with max -log10(padj) value shown in parentheses in motif label. P-values were calculated via hypergeometric test in ArchR^32^. **d.** UMAP colored by *C1QB* normalized gene expression (**left**). *C1QB* locus (chr1:22,652,235-22,653,595) with selected isoforms, motifs, open chromatin peaks, and chromatin accessibility reads from unimodal and multimodal ATAC cells aggregated by chromatin class and scaled by read counts per class (**Methods**) (**right**).

We next investigated the TF motifs enriched in the myeloid chromatin classes. M_A_-2 was enriched for KLF motifs (**Fig. 4c**), with *KLF4* (p_adj_=1.34e-6) known to both establish residency of TRMs and to assist in their phagocytic function^61^. Furthermore, we found a KLF4 motif in the promoter of *C1QB*, whose protein product bridges phagocytes to the apoptotic cells they clear^62^ (**Fig. 4d**). Both the intermediate M_A_-4 and the infiltrating monocyte M_A_-1 classes had significant enrichments of AP-1 activation motifs^63^ (JUN p_adj_=1.77e-153, 3.65e-136, respectively; **Fig. 4c**). AP-1 factors have been shown to function in human classical monocytes along with CEBP factors^64^, also enriched in M_A_-1 (CEBPD p_adj_=2.10e-26; **Fig. 4c**). SPI1 (PU.1) is the master regulator of myeloid development^65^, including conventional DCs^66^. We found PU.1 motifs most strongly enriched in the DC cluster M_A_-3 (p_adj_=3.24e-55; **Fig. 4c**).

### RA B/plasma chromatin classes

Next, we clustered 8,641 B and plasma cells into 4 MS4A1*+* B cell and 2 SDC1*+* plasma cell chromatin classes (**Methods**; **Fig. 5a**; **Supplementary Fig. 5a**). We defined a B_A_-3: FCER2+ IGHD+ naive B class with high accessibility and expression of *FCER2* encoding naïve marker CD23^67^ (**Fig. 5b**; **Supplementary Fig. 5b**). We also labeled a B_A_-4: CD24+ MAST4+ unswitched memory B class (**Supplementary Fig. 5b**). *IGHD* and *IGHM* expression was lower in B_A_-2: TOX+ PDE4D+ switched memory B cells, and the TF TOX had its highest expression and accessibility within B cells in B_A_-2 as previously shown in switched memory B cells^68, 69^ (**Fig. 5b**; **Supplementary Fig. 5b**). B_A_-5: ITGAX+ ABC (Age-Associated B cells) had high accessibility and expression of *ITGAX*, which encodes for CD11c, a key ABC marker^70^ (**Fig. 5b**; **Supplementary Fig. 5b**). ABCs were shown to be associated with leukocyte-rich RA^11^ with a potential role in antigen presentation^71^, which was supported here by expression of LAMP1 and HLA-DRA in B_A_-5 (**Supplementary Fig. 5b**). The plasma chromatin class, B_A_-0: CREB3L2+ plasma, was marked by the TF *CREB3L2*, a known factor in the transition between B and plasma cells^72^ (**Fig. 5B**; **Supplementary Fig. 5b**). These results suggested tissue *in situ* B cell activation and differentiation into plasma cells, as we have previously suggested^73^. Finally, B_A_-1: CD27+ plasma, had the highest accessibility and expression of *CD27* (**Fig. 5b**; **Supplementary Fig. 5b**). We note that plasma cells were difficult to define using ATAC data, with many of the immunoglobulin genes having a paucity of chromatin accessibility (**Supplementary Fig. 5b**).

We then explored the TF motif landscape of B and plasma cells. B cells shared many TF motifs across clusters, with many ETS factors (*e.g.*, SPIB, SPI1, ETS1) as well as EBF1 and NFkB1/2 (**Fig. 5c**). SPIB and SPI1 work together to regulate B cell receptor signaling^74^, which starts its dysregulation in RA at the naive B cell level^75, 76^ (p_adj_=0, 0, respectively; **Fig. 5c**). Switched memory B cells were enriched for ETS1 motifs (p_adj_=9.51e-19; **Fig. 5c**), whose TF is required for IgG2a class switching in mice^77^. In plasma cells, B_A_-0 had motifs such as KLF2^78^ and SP3^79^ (p_adj_=8.94e-105, 3.84e-138, respectively; **Fig. 5c-d**). B_A_-1 was enriched for AP-1 factor motifs^80^, namely BATF::JUN (p_adj_=0; **Fig. 5c-d, Supplementary Fig. 5c**). In the locus of PRDM1, a known plasma TF^79^, the more B_A_-0 accessible peak had an SP3 motif while the more B_A_-1 accessible peaks had BATF::JUN motifs (**Fig. 5d**), suggesting potentially different regulatory strategies by class.

**Fig. 5.**
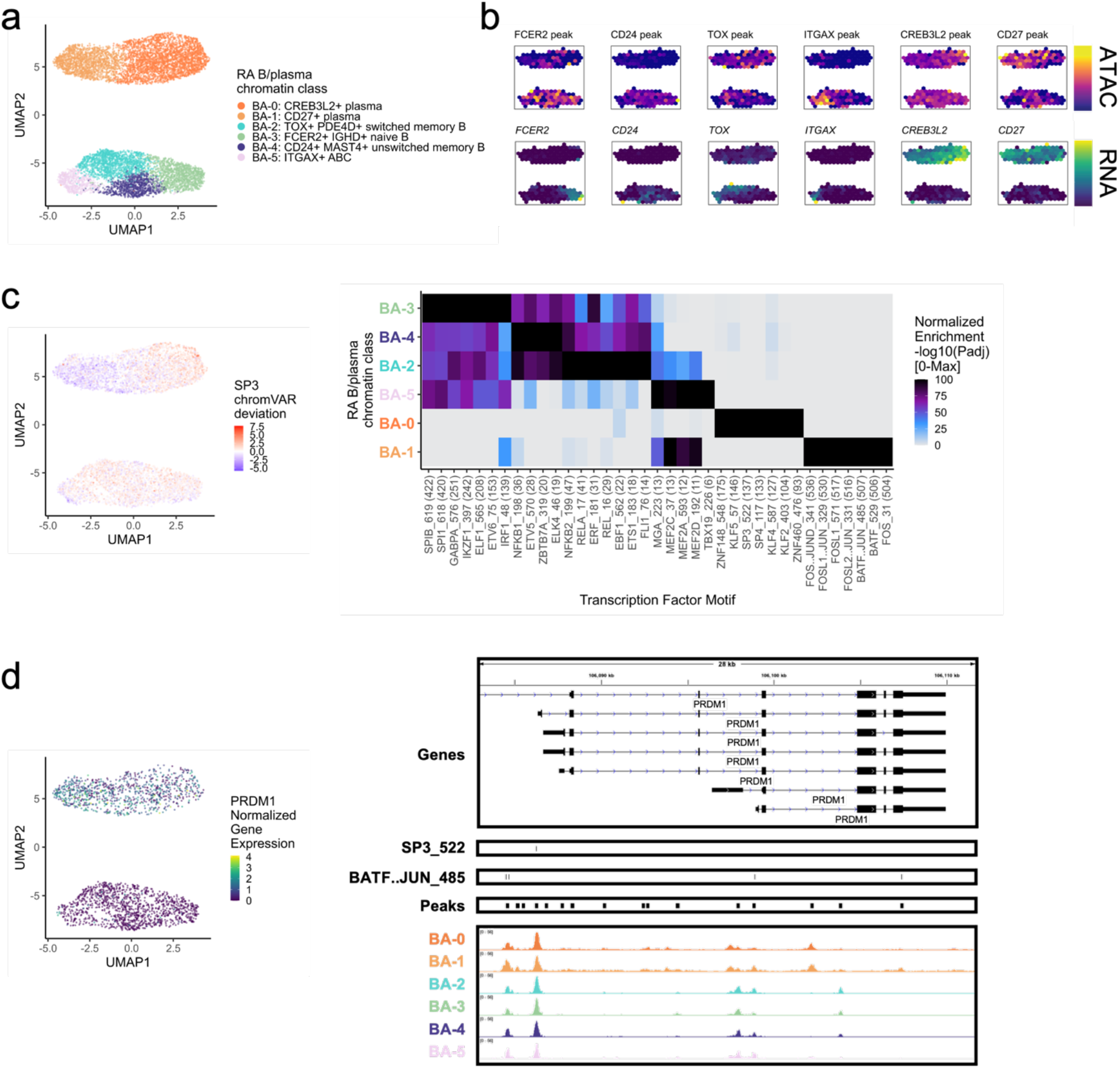
RA B/plasma chromatin classes. **a.** UMAP colored by 6 B/plasma chromatin classes defined from unimodal scATAC and multimodal snATAC cells. **b.** Binned normalized marker peak accessibility (**top**) and gene expression (**bottom**) for multiome snATAC cells on UMAP. **c.** UMAP colored by chromVAR^31^ deviations for the SP3 motif (**left**). Most significantly enriched motifs in marker peaks per B/plasma chromatin class (**right**). To be included per class, motifs had to be enriched in the class above a minimal threshold and corresponding TFs had to have at least minimal expression in snRNA (**Methods**). Color scale normalized per motif across classes with max -log10(padj) value shown in parentheses in motif label. P-values were calculated via hypergeometric test in ArchR^32^. **a. d.** UMAP colored by *PRDM1* normalized gene expression (**left**). *PRDM1* locus (chr6:106,082,865-106,111,658) with selected isoforms, motifs, open chromatin peaks, and chromatin accessibility reads from unimodal and multimodal ATAC cells aggregated by chromatin class and scaled by read counts per class (**Methods**) (**right**).

### RA endothelial chromatin classes

Among the 3,809 endothelial cells, we identified 4 chromatin classes (**Fig. 6a**; **Supplementary Fig. 6a**). The E_A_-2: SEMA3G+ arteriolar class had gene and peak markers for signaling-related genes including *SEMA3G*^81^, *CXCL12*, and *JAG1* (**Fig. 6b**; **Supplementary Fig. 6b**). The NOTCH3 signaling gradient that causes inflammation and joint destruction in RA mouse models likely originates through Notch ligand *JAG1* in these arteriolar endothelial cells^21^. We identified the E_A_-0: SELP+ venular class with markers for leukocyte trafficking to tissue such as *SELP*^82^ as well as inflammatory genes *HLA-DRA* and *CD74* (**Fig. 6b**; **Supplementary Fig. 6b**). We also found a capillary class, E_A_-1: RGCC+ capillary marked by *RGCC* and *SPARC*^83^ chromatin accessibility and gene expression (**Fig. 6b**; **Supplementary Fig. 6b**). Finally, a small population of E_A_-3: PROX1+ lymphatic cells had gene expression of and promoter accessibility at *PROX1*^84^ and *PARD6G* genes (**Fig. 6b**; **Supplementary Fig. 6b**).

**Fig. 6.**
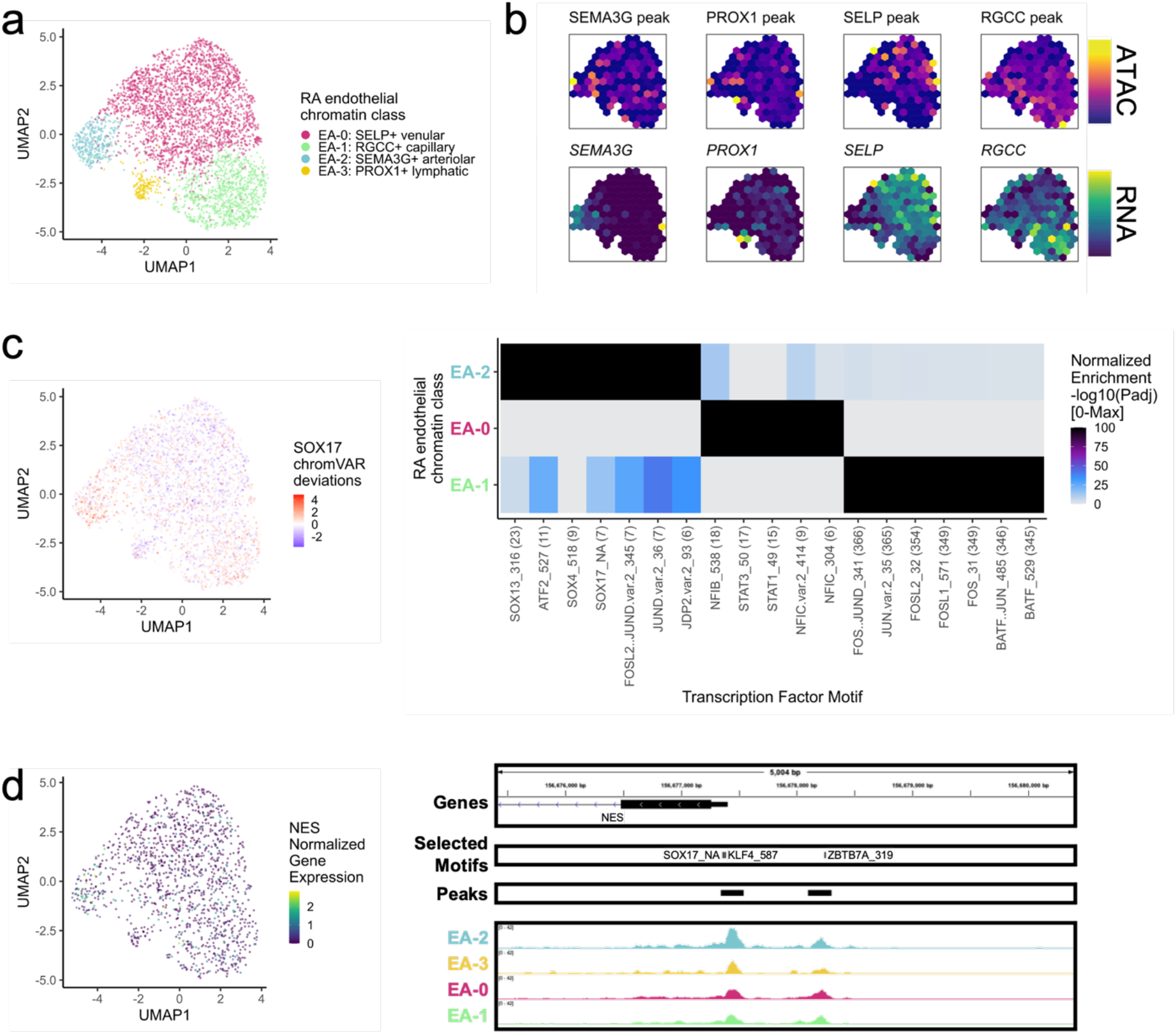
RA endothelial chromatin classes. **a.** UMAP colored by 4 endothelial chromatin classes defined from unimodal scATAC and multimodal snATAC cells. **b.** Binned normalized marker peak accessibility (**top**) and gene expression (**bottom**) for multiome snATAC cells on UMAP. **c.** UMAP colored by chromVAR^31^ deviations for the SOX17 motif (**left**). Most significantly enriched motifs in marker peaks per endothelial chromatin class (**right**). To be included per class, motifs had to be enriched in the class above a minimal threshold and corresponding TFs had to have at least minimal expression in snRNA (**Methods**). Color scale normalized per motif across classes with max -log10(padj) value shown in parentheses in motif label. P-values were calculated via hypergeometric test in ArchR^32^. E_A_-3 is not shown because only 1 marker peak was found, likely due to low cell counts. **d.** UMAP colored by *NES* normalized gene expression (**left**). *NES* locus (chr1:156,675,399-156,680,400) with selected isoforms, motifs, open chromatin peaks, and chromatin accessibility reads from unimodal and multimodal ATAC cells aggregated by chromatin class and scaled by read counts per class (**Methods**) (**right**).

We identified SOX motifs^85^ in E_A_-2, STAT motifs^86^ in E_A_-0, and AP-1 motifs^87^ in E_A_-1 (**Fig. 6c**). *Sox17* is a crucial intermediary between Wnt and Notch signaling that specifically initiates and maintains endothelial arterial identity in mice^85^. Similarly, we found a SOX17 motif (p_adj_=3.27e-8) in the promoter of *NES*^88, 89^ with its highest accessibility and expression in E_A_-2 cells (**Fig. 6d**).

### Synovial tissue is key to identifying pathogenic RA chromatin classes

To determine if the chromatin classes identified in RA tissue were comparable with the known peripheral blood chromatin landscape, we clustered the tissue cells with those from a published healthy PBMC multiome dataset^90, 91^ (**Methods**; **Supplementary Fig. 7**). To determine the similarity between the PBMC and tissue chromatin classes, we calculated the Odds Ratio (OR) between the newly defined clusters and the previous blood and tissue labels; overall, there was good concordance. For example, the PBMC Treg cells and T_A_-3: CD4+ IKZF2+ Treg cells were grouped in combined cluster 5 (OR: 12 and 85, respectively) (**Supplementary Fig. 7a**) and PBMC cDC1, cDC2, and pDCs all associated with M_A_-3: CD1C+ AFF3+ DCs in combined cluster 4 (OR: Infinite, 45, 78, and 98, respectively) (**Supplementary Fig. 7b**). However, there were some tissue chromatin classes that did not have clear counterparts in PBMCs, such as T_A_-2: CD4+ PD-1+ TFH/TPH, M_A_-2: LYVE1+ TIMD4+ TRM, M_A_-4: SPP1+ FABP5+ intermediate, and B_A_-5: ITGAX+ ABC (**Supplementary Fig. 7**). Intriguingly, these chromatin classes only identified in the RA synovial tissue are known to be important in RA pathogenesis^11, 13, 14, 16, 60^. While this could be a difference between healthy and disease states beyond the blood and tissue comparison, these populations generally skew towards tissue populations^13, 92, 93^ and suggested the importance of examining cells from diseased tissue environments.

### Chromatin classes are epigenetic superstates of transcriptional cell states

To understand how these chromatin classes corresponded to transcriptionally defined cell states, we used Symphony^94^ to map the RA multimodal snRNA profiles into the well-annotated AMP-RA cell type references^12^. After embedding the multimodal snRNA profiles into the AMP-RA reference data, we annotated each multimodal cell by the most common cell state of its five nearest reference neighbors (**Methods**). 70% of T cells (24 states), 96% of stromal cells (10 states), 96% of myeloid cells (15 states), 96% of B/plasma cells (9 states), and 99% of endothelial cells (5 states) mapped well (*i.e.*, 3/5 neighbors had the same cell state annotation). We also observed that the proportion of each cell state in the AMP-RA reference and the multimodal query datasets was consistent, suggesting that the reference and query datasets have comparable cell state distributions despite different technologies (**Supplementary Fig. 8a-e**).

We then sought to understand the correspondence between the mapped transcriptional cell states and chromatin classes. We calculated an OR for each combination of state and class to measure the strength of association and used a Fisher’s exact test to assess significance (**Methods**).

We observed that each transcriptional cell state generally corresponded to a single chromatin class (**Fig. 7**; **Supplementary Figure S8g-h**). In contrast, a single chromatin class represents a superstate encompassing multiple transcriptionally defined cell states. For example, cells in the T_A_-0: CD8+ GZMK+ chromatin class were more likely to be labelled in the T-5: CD4+ GZMK+ memory, T-13: CD8+ GZMK/B+ memory, and T-14: CD8+ GZMK+transcriptional cell states across CD4/CD8 lineages (OR=11, 12, 11, respectively; **Fig. 7a**); the high *GZMK* promoter accessibility and expression shared by these states may contribute to this categorization (**Supplementary Fig. 8f**). We saw examples of this model in every cell type: S_A_-1 linked to F-0/F-1 and S_A_-0 to F-6/F-5/F-3/F-8 in stromal cells; M_A_-1 to M-7/M-11 and M_A_-4 to M-3/M-4 in myeloid cells; B_A_-4 to B-1/B-3 in B/plasma cells; and E_A_-0 to E-1/E-2 in endothelial cells as more examples (**Fig. 7b-c**; **Supplementary Figure S8g-h**; **Supplementary Table 4**). Indeed, when we aggregated the snATAC reads by states, we observed shared openness between transcriptional cell states within the same class (*i.e.*, superstate), as seen with the cytotoxic T_A_-4 grouped cell states T-12/T-15 at the cytotoxicity-associated^34^ *FGFBP2* gene, lining fibroblast S_A_-1 grouped cell states F-0/F-1 at the lining-associated^11^ *CLIC5* gene, and intermediary myeloid M_A_-4 grouped cell states M-3/M-4 at bone marrow macrophage-associated^59^ *SPP1* gene (**Supplementary Fig. 9**).

**Fig. 7.**
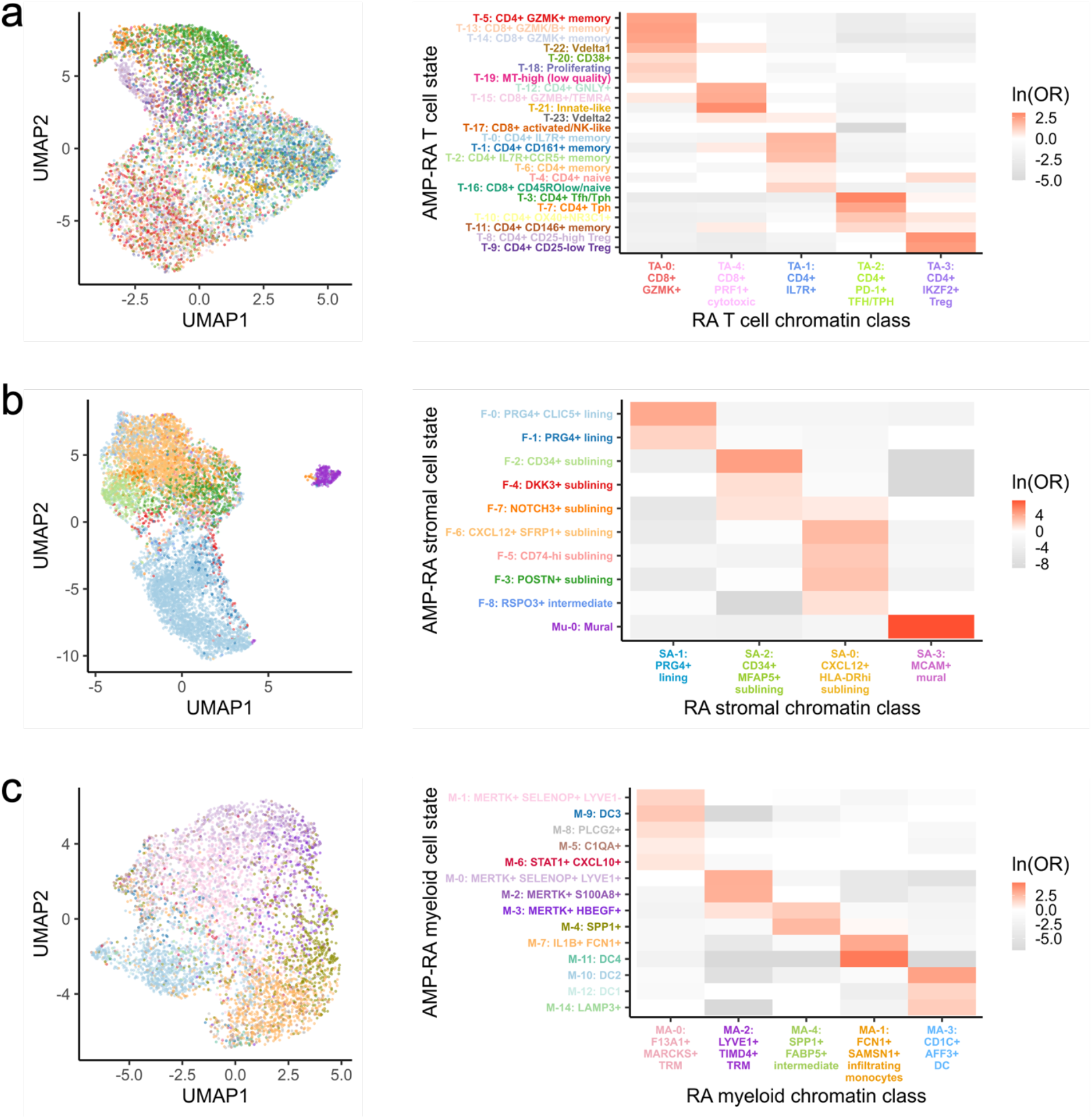
A chromatin class encompassed multiple transcriptional cell states in proposed superstate model. For (**a.**) T, (**b.**) stromal, and (**c.**) myeloid cells, UMAP colored by classified AMP-RA reference transcriptional cell states for multiome cells (**left**) and natural log of Odds Ratio between chromatin classes and transcriptional cell states (**right**). Non-significant values (FDR<0.05) are white. In **c.**, M-13: pDC transcriptional cell state was excluded as fewer than 10 cells were classified into it.

We next asked if evidence for chromatin superstates was sensitive to clustering resolution. We observed that the class and state relationships largely replicated when we increased the open chromatin clustering resolution (**Supplementary Fig. 10**). To further support the superstate hypothesis, we trained a linear discriminant analysis (LDA) model to predict the transcriptional cell state between each pair of states from the ATAC principal components (PCs), upon which the chromatin classes were defined (**Methods**). Generally, transcriptional cell states belonging to the same chromatin class were difficult to distinguish using ATAC data alone (**Supplementary Fig. 11**). For example, transcriptional states T-14 and T-13 both belonged to chromatin class T_A_-0, and thus ATAC PCs could not easily discriminate between them (AUROC=0.61); on the other hand, T-14 and T-3 belonged to classes T_A_-0 and T_A_-2, respectively, and LDA nearly perfectly distinguished them (AUROC=0.98) (**Supplementary Fig. 11a**). In all cell types, the mean AUROC between states within the same chromatin class was less than that of states across different chromatin classes. For example in T cells, the mean AUROC was 0.77 within the same classes and 0.88 across different chromatin classes, suggesting that there was a limit to how well the ATAC data could differentiate between transcriptional cell states.

### Cell neighborhood associations with histological metrics and cell state proportions

Next, we sought to investigate associations between the RA chromatin classes and RA clinical metrics using the larger AMP-RA reference dataset with clinical measurements for 79 RA or OA patients. Per cell type, we classified^94^ each cell from the AMP-RA reference dataset, now the query, into the RA chromatin classes based on the five nearest multiome snRNA neighbors, now the reference (**Methods**). To validate this annotation, we compared the relative proportions of chromatin classes between the unimodal scATAC cells and the projected AMP-RA scRNA cells for donors in both studies (**Methods**). We observed generally high correlation between the two technologies (**Fig. 8a**; **Supplementary Fig. 12a**). We then investigated RA clinical associations calculated via Co-varying Neighborhood Analysis (CNA)^95^. In brief, CNA tests associations between sample-level attributes, such as clinical metrics, and cellular neighborhoods, which are small groups of cells that reflect granular cell states. We used the previously described CNA associations defined in the AMP-RA reference cells and re-aggregated them by their chromatin classes (**Methods**). For example, we found an association between myeloid cells and histology characterized by lymphoid infiltration density (p=0.005).

**Fig. 8.**
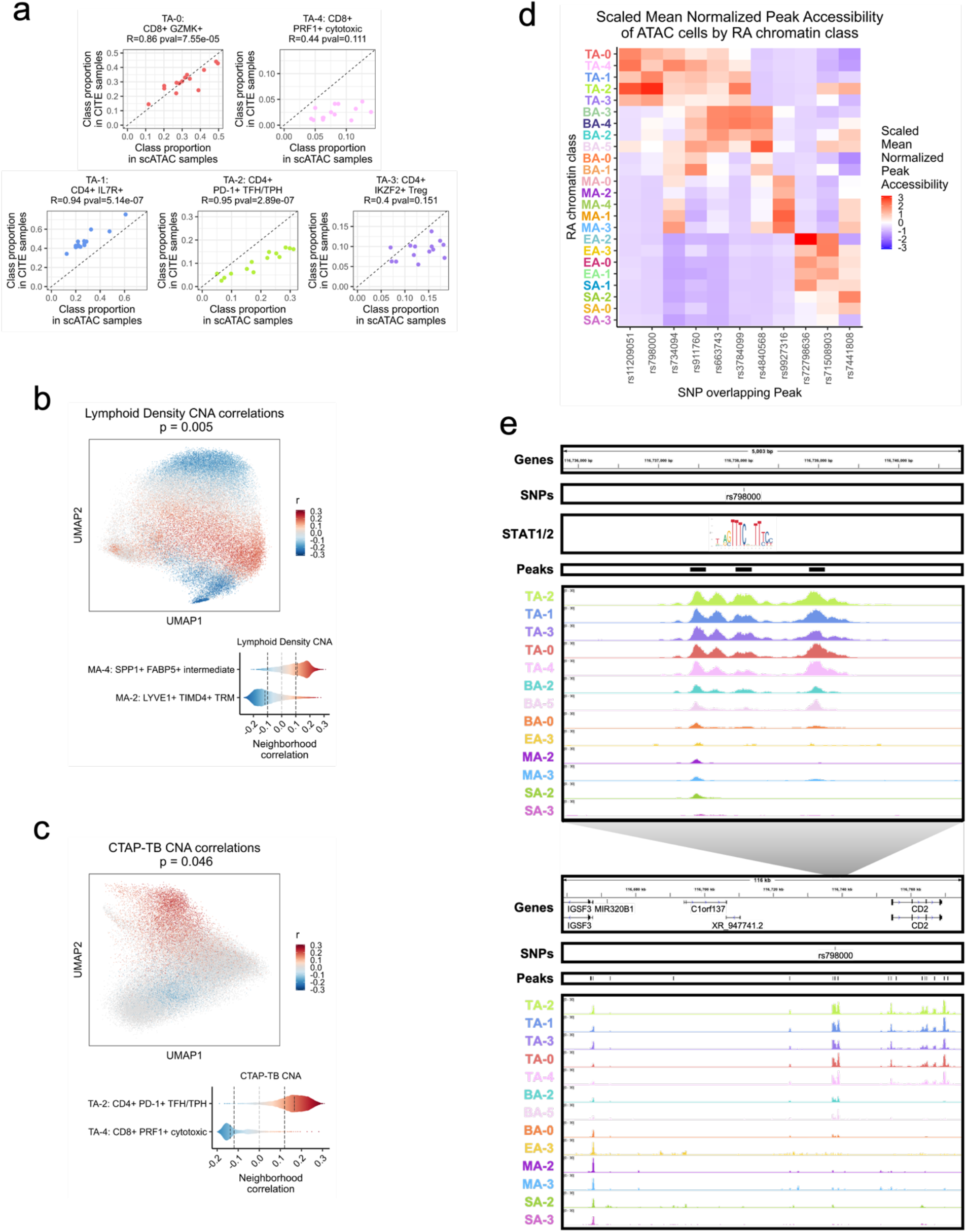
Linking RA chromatin classes to RA pathology. **a.** For each donor shared between the unimodal ATAC and AMP-RA reference studies with at least 200 T cells, the Pearson correlation between the relative proportions of T cell chromatin classes defined in the unimodal ATAC datasets (**x-axis**) and classified into in the CITE datasets through the multiome cells (**y-axis**). Pearson Correlation Coefficients (R) and p-values (pval) noted. **b.** CNA correlations between myeloid cell neighborhoods and lymphoid density in AMP-RA reference myeloid cells visualized on UMAP (**top**) and aggregated by classified myeloid chromatin classes (**bottom**). On the top, cells not passing the FDR threshold were colored grey. On the bottom, FDR thresholds shown in dotted black lines. **c.** CNA correlations between T cell neighborhoods and CTAP-TB in AMP-RA reference T cells visualized on UMAP (**top**) and aggregated by classified T cell chromatin classes (**bottom**). On the top, cells not passing the FDR threshold were colored grey. On the bottom, FDR thresholds shown in dotted black lines. **d.** Scaled mean normalized chromatin accessibility for peaks that overlap putatively causal RA risk variants across chromatin classes. Additional information in **Supplementary Table 5**. **e.** rs798000 locus, zoomed in (chr1:116,735,799-116,740,800) (**top**) and zoomed out (chr1:116,658,581-116,775,106) (**bottom**) with isoforms, SNPs, open chromatin peaks, and chromatin accessibility reads aggregated by chromatin class and scaled by read counts per class (**Methods**). STAT1/2 motif was downloaded from JASPAR^98^ ID MA0517.1 and is not to scale, but it is aligned to the SNP-breaking motif position.

Specifically, the increase in lymphocyte populations was positively associated with M_A_-4: SPP1+ FABP5+ intermediate class, whose inflammatory cytokines/chemokines production may be responsible for lymphocyte homing^96^, and negatively associated with M_A_-2: LYVE1+ TIMD4+ TRM, whose gene markers were found more often on synovial TRMs from healthy and remission RA than active RA patients^14^ (**Fig. 8b**). Additionally, we observed an association between T cells and histological Krenn inflammation score (p=0.02), with T_A_-2: CD4+ PD-1+ TFH/TPH positively^97^ and T_A_-4: CD8+ PRF1+ cytotoxic negatively correlated (**Supplementary Fig. 12b**). These results were consistent with the original transcriptional cell state findings^12^ and suggested that the connections between RA pathology and cell state may begin before transcription.

One of the key findings from the AMP-RA study was the identification of six Cell Type Abundance Phenotypes (CTAPs), which characterized RA patients into subtypes based on the relative proportions of their broad cell type abundances in synovial tissue^12^. For example, CTAP-TB has primarily T and B/plasma cells. Specific cell neighborhoods within cell types were expanded or depleted in these CTAPs as defined by CNA associations in the AMP-RA reference cells. We recapitulated some of these transcriptional associations by re-aggregating the CNA results within the chromatin classes; for example, the RA T cell class T_A_-2 was positively associated with CTAP-TB compared to other T cell states, likely reflecting the role of TFH/TPH cells in B cell inflammation response^11, 13^, while T_A_-4 was negatively associated (p=0.046; **Fig. 8c**). Furthermore, in stromal cells, we saw the S_A_-1: PRG4+ lining class positively associated with CTAP-F, a primarily fibroblast CTAP (p=0.0027; **Supplementary Fig. 12c**). This suggested that the most expanded type of fibroblasts in CTAP-F individuals was predominantly from the synovial lining layer, which was consistent with lining marker CLIC5 protein having high staining in the lining fibroblasts and being expressed in the highest proportion of cells from high density fragments of CTAP-F samples (ANOVA p_adj_<0.001 between CTAPs)^12^. Therefore, we could meaningfully replicate the RA pathological associations of both clinical metrics and phenotypic subtypes to transcriptional cell states using their related chromatin class superstate, suggesting that the epigenetic regulation underlying the transcriptional cell states may be mined for further pathological insights into RA.

### Chromatin classes prioritize RA-associated SNPs

We next asked whether RA risk variants overlap the chromatin classes to help define function for putatively causal variants, genes, and pathways at play in RA pathology^99–103^. Using an RA multi-ancestry genome-wide association meta-analysis study^104^, we overlapped fine-mapped non-coding variants with posterior inclusion probability (PIP) greater than 0.1 with the 200 bp open chromatin peaks and assessed peak accessibility across the 24 chromatin classes (**Methods**; **Fig. 8D; Supplementary Table 5**). For six loci, putatively causal variants overlapped a peak accessible in predominantly one cell type, such as rs11209051 in peak chr1:67333106-67333306 in T cells (Wilcoxon T versus non-T class one-sided p=4.17e-04; **Methods**) near the *IL12RB2* gene and rs4840568 in peak chr8:11493501-11493701 in B/plasma cells (Wilcoxon p=1.49e-05) near the *BLK* gene. In the other loci, variants overlapped with chromatin classes from 2 cell types, with most combinations involving T cells. Moreover, there were 4 SNPs overlapping peaks accessible in the T_A_-2: CD4+ PD-1+ TFH/TPH class, which is the most targeted class within T cells and important for RA pathogenesis^11, 13^.

As an example, we observed putatively causal SNP rs798000 (PIP=1.00) overlap peak chr1:116737968-116738168, accessible primarily in T cells (Wilcoxon p=2.35e-05) with T_A_-2 as its most accessible class (z=3.03) (**Fig. 8d-e**, top). In a previous study^91^, we linked active chromatin regions to their target genes, which suggested *CD2* is a causal gene in this locus. *CD2* is a co-stimulatory receptor primarily expressed on T and NK cells^105^, which likely explains why it was only accessible in our T cell chromatin classes among the five cell types investigated (**Fig. 8e**, bottom). Intriguingly, rs798000 overlaps a STAT1/2 binding site at a high information content half site position (**Fig. 8e**, top, position 8 in JASPAR^98^ motif MA0517.1), suggesting a potential direct link to TF regulation of the JAK/STAT pathway commonly upregulated in RA^52^.

We also discovered SNP rs9927316 (PIP=0.54) in myeloid-specific peak chr16:85982638-85982838 (Wilcoxon p=4.165e-04), downstream of *IRF8*, one of the master regulator TFs of myeloid and B cell fates^106–108^ (**Supplementary Fig. 13a**). The SNP disrupts a KLF4 motif^61^, one of the TRM TFs highlighted earlier (**Supplementary Fig. 13a**; **Fig. 4c-d**). Furthermore, we observed SNP rs734094 (PIP=0.41) overlapping peak chr11:2301916-2302116, with its most accessible classes in T and myeloid cells: T_A_-4: CD8+ PRF1+ cytotoxic and M_A_-3: CD1C+ AFF3+ DC (z=1.94, 1.65, respectively) (**Fig. 8d**; **Supplementary Fig. 13b**). While existing in the promoters of both *TSPAN32* and *C11orf21* gene isoforms (**Supplementary Fig. 13b**), we^91^ proposed the causal gene as Lymphocyte-specific Protein 1 (*LSP1*), shown to negatively regulate T cell migration and T cell-dependent inflammation in arthritic mouse models^109^. For each of these loci, we also aggregated chromatin accessibility by classified transcriptional cell state and saw that the multiple states underlying each class had similar patterns, such as rs734094 having some of the strongest signal in T_A_-4 associated classes T-12, T-21 and M_A_-3 associated classes M-10, M-14 (**Supplementary Fig. 14**). This both reaffirmed our chromatin class superstate model and suggested that the classes are useful functional units that may help simplify mapping risk loci to affected cell states. The RA tissue chromatin classes can help prioritize putative cell states of action for non-coding RA risk variants that may help assist in their functional characterization within disease etiology.

## Discussion

In this study, we described 24 chromatin classes across 5 broad cell types in 30 synovial tissue samples assayed with unimodal scATAC and multimodal snATAC along with TFs potentially regulating them. Based on our observation that cells from the same chromatin class corresponded to multiple transcriptional cell states, we proposed that these chromatin classes are putative superstates of related transcriptional cell states. Finally, we assessed these chromatin classes’ relationship to RA clinical metrics, subtypes, and genetic risk variants.

Simultaneous chromatin accessibility and gene expression measurements in the multiome cells were essential to test the relationship between chromatin classes and transcriptional cell states. Biologically, open chromatin is necessary but not sufficient for gene expression^18^, so it is reasonable to expect related cell states to have similar open chromatin landscapes with poised enhancers activated by specific TFs in the required state. The robustness of the observed class-state relationships across multiple clustering resolutions mitigated concerns that this proposed model was a technical artifact. Moreover, even in the absence of clusters, classifiers based on continuous ATAC PCs also demonstrated the similarity of transcriptional states within the same chromatin class.

Defining the relationship between transcriptional cell state and chromatin class may have important therapeutic implications. One effective RA treatment strategy is the deletion of the pathogenic cell state: the use of B-cell depleting antibodies (*e.g.*, rituximab^10^) is an example. However, if one chromatin class corresponds to multiple transcriptional cell states, then deleting very specific pathogenic populations may be ineffective as other non-pathogenic transcriptional cell states may transition into the specific pathogenic cell state in response to the same pathogenic tissue environment. In that case, altering the environment or removing exogenous factors *(e.g.*, TFs, cytokines) might be a more effective treatment. S_A_-0: CXCL12+ HLA-DR^hi^ sublining fibroblasts, with its four related transcriptional states in our superstate model, may be an interesting class to study in this regard. S_A_-0 accessible peaks were enriched for STAT motifs, suggesting potential regulation by the JAK/STAT signaling pathway. Indeed, JAK inhibition via tofacitinib and upadacitinib has been shown to prevent HLA-DR induction in RA synovial fibroblasts^110^.

More broadly, the results presented here suggest some interesting next steps. First, our chromatin class superstate model indicated that certain transcriptional cell states were more closely linked, but further experimentation would be required to ascertain whether these related cell states have a plastic enough chromatin landscape that they can potentially cross-differentiate within a cell type or whether they are more broadly grouped by function. Second, to better understand whether the more pathogenic chromatin classes such as T_A_-2: CD4+ PD-1+ TFH/TPH and M_A_-1: FCN1+ SAMSN1+ infiltrating monocytes are indeed only in tissue, a RA PBMC scATAC-seq study may be warranted. If we see more of a consensus between the chromatin landscapes of RA blood and tissue, we may be able to determine if the chromatin environment is permissible for some of these pathogenic transcriptional populations to arise before they do. If not, then we confirm the need to investigate tissue inflammation directly at the tissue level. Third, the chromatin classes could prioritize where to look for functional effects of putatively causal RA genetic variants. For example, further study could investigate whether STAT signaling upon *CD2* stimulation^111, 112^ is affected by the STAT1/2-motif breaking SNP rs798000 in TFH/TPH cells, in particular from donors with a subtype of RA characterized by primarily T and B/plasma cells, as in CTAP-TB, where TFH/TPH cells are most positively correlated. Our study underscores the value for larger tissue-specific genetic studies examining the role of genetic variation on open chromatin.

In conclusion, we presented an atlas for RA tissue chromatin classes that will be a useful resource for linking chromatin accessibility to gene expression and the interpretation of genetic information.

## Accelerating Medicines Partnership Program: Rheumatoid Arthritis and Systemic Lupus Erythematosus (AMP RA/SLE) Network includes

Jennifer Albrecht^7^, William Apruzzese^11^, Nirmal Banda^18^, Jennifer L. Barnas^7^, Joan M. Bathon^12^, Ami Ben-Artzi^13^, Brendan F. Boyce^14^, David L. Boyle^15^, S. Louis Bridges Jr.^8, 9^, Vivian P. Bykerk^8,9^, Debbie Campbell^7^, Hayley L. Carr^16^, Arnold Ceponis^15^, Adam Chicoine^1^, Andrew Cordle^17^, Michelle Curtis^1,2,3,4,5^, Kevin D. Deane^18^, Edward DiCarlo^19^, Patrick Dunn^20, 21^, Andrew Filer^16^, Gary S. Firestein^15^, Lindsy Forbess^16^, Laura Geraldino-Pardilla^12^, Susan M. Goodman^8,9^, Ellen M. Gravallese^1^, Peter K. Gregersen^22^, Joel M. Guthridge^23^, Maria Gutierrez-Arcelus^1,2,3,4,5,24^, Siddarth Gurajala^1,2,3,4,5^, V. Michael Holers^18^, Diane Horowitz^22^, Laura B. Hughes^25^, Kazuyoshi Ishigaki^1,2,3,4,5,26^, Lionel B. Ivashkiv^8,9^, Judith A. James^23^, Joyce B. Kang^1,2,3,4,5^, Gregory Keras^1^, Ilya Korsunsky^1,2,3,4,5^, Amit Lakhanpal^8,9^, James A. Lederer^27^, Zhihan J. Li^1^, Yuhong Li^1^, Katherine P. Liao^1,4^, Arthur M. Mandelin II^28^, Ian Mantel^8,9^, Mark Maybury^16^, Andrew McDavid^29^, Joseph Mears^1,2,3,4,5^, Nida Meednu^7^, Nghia Millard^1,2,3,4,5^, Larry W. Moreland^18,30^, Alessandra Nerviani^31^, Dana E. Orange^8,32^, Harris Perlman^28^, Costantino Pitzalis^31^, Javier Rangel-Moreno^7^, Karim Raza^16^, Yakir Reshef^1,2,3,4,5^, Christopher Ritchlin^7^, Felice Rivellese^31^, William H. Robinson^33^, Laurie Rumker^1,2,3,4,5^, Ilfita Sahbudin^16^, Jennifer A. Seifert^18^, Kamil Slowikowski^4,5,34,35^, Melanie H. Smith^8^, Darren Tabechian^7^, Dagmar Scheel-Toellner^16^, Paul J. Utz^33^, Dana Weisenfeld^1^, Michael H. Weisman^13,33^, Qian Xiao^1,2,3,4,5^

^11^ Accelerating Medicines Partnership® Program: Rheumatoid Arthritis and Systemic Lupus Erythematosus (AMP® RA/SLE) Network

^12^ Division of Rheumatology, Columbia University College of Physicians and Surgeons, New York, NY, USA.

^13^ Division of Rheumatology, Cedars-Sinai Medical Center, Los Angeles, CA, USA.

^14^ Department of Pathology and Laboratory Medicine, University of Rochester Medical Center, Rochester, NY, USA

^15^ Division of Rheumatology, Allergy and Immunology, University of California, San Diego, La Jolla, CA, USA.

^16^ Rheumatology Research Group, Institute for Inflammation and Ageing, University of Birmingham, NIHR Birmingham Biomedical Research Center and Clinical Research Facility, University of Birmingham, Queen Elizabeth Hospital, Birmingham, UK.

^17^ Department of Radiology, University of Pittsburgh Medical Center, Pittsburgh, PA, USA.

^18^ Division of Rheumatology, University of Colorado School of Medicine, Aurora, CO, USA.

^19^ Department of Pathology and Laboratory Medicine, Hospital for Special Surgery; New York, NY, USA.

^20^ Division of Allergy, Immunology, and Transplantation, National Institute of Allergy and Infectious Diseases, National Institutes of Health, Bethesda, MD, USA.

^21^ Northrop Grumman Health Solutions, Rockville, MD, USA.

^22^ Feinstein Institute for Medical Research, Northwell Health, Manhasset, New York, NY, USA.

^23^ Department of Arthritis & Clinical Immunology, Oklahoma Medical Research Foundation, Oklahoma City, OK, USA.

^24^ Division of Immunology, Department of Pediatrics, Boston Children’s Hospital and Harvard Medical School, Boston, MA. US.

^25^ Division of Clinical Immunology and Rheumatology, Department of Medicine, University of Alabama at Birmingham, Birmingham, AL, USA.

^26^ Laboratory for Human Immunogenetics, RIKEN Center for Integrative Medical Sciences, Yokohama, Japan.

^27^ Department of Surgery, Brigham and Women’s Hospital and Harvard Medical School, Boston, MA, USA.

^28^ Division of Rheumatology, Department of Medicine, Northwestern University Feinberg School of Medicine, Chicago, IL, USA.

^29^ Department of Biostatistics and Computational Biology, University of Rochester School of Medicine and Dentistry; Rochester, NY, USA.

^30^ Division of Rheumatology and Clinical Immunology, University of Pittsburgh School of Medicine; Pittsburgh, PA, USA.

^31^ Centre for Experimental Medicine & Rheumatology, William Harvey Research Institute, Queen Mary University of London; London, UK.

^32^ Laboratory of Molecular Neuro-Oncology, The Rockefeller University, New York, NY, USA.

^33^ Division of Immunology and Rheumatology, Institute for Immunity, Transplantation and Infection, Stanford University School of Medicine, Stanford, CA, USA.

^34^ Center for Immunology and Inflammatory Diseases, Department of Medicine, Massachusetts General Hospital (MGH), Boston, MA, USA

^35^ MGH Cancer Center, Boston, MA, USA

## Methods

### Patient recruitment

Fourteen RA and 4 OA patients were recruited by the Accelerating Medicines Partnership (AMP) Network for RA and SLE to provide samples for use in the unimodal scATAC-seq experiments. Separately, synovial tissue samples from 11 RA patients and 1 OA patient were collected from Brigham and Women’s Hospital (BWH) and the Hospital for Special Surgery (HSS) for use in the multimodal ATAC + Gene Expression experiments. Histologic sections of RA synovial tissue were examined, and samples with inflammatory features were selected in both cases.

All clinical and experimental sites that recruited patients obtained approval for this study from their Institutional Review Boards. All patients gave informed consent. We have complied with all relevant ethical regulations.

### Synovial tissue collection and preparation

Synovial tissue samples from 14 RA patients and 4 OA patients were collected and cryopreserved as part of a larger study cohort by the AMP Network for RA and SLE, as previously described^12^. Synovial tissue samples were thawed and disaggregated as previously described^12,23^. The resulting single-cell suspensions were stained with anti-CD235a antibodies (clone 11E4B-7-6 (KC16), Beckman Coulter) and Fixable Viability Dye (FVD) eFlour 780 (eBioscience/ThermoFisher). Live non-erythrocyte (*i.e.*, FVD-CD235-) cells were collected by fluorescence-activated cell sorting (BD FACSAria Fusion). The sorted live cells were then re-frozen in Cryostor and stored in liquid nitrogen. The cells were later thawed and processed as described above for droplet-based scATAC-seq according to manufacturer’s protocols (10X Genomics). For the multimodal experiments, the 11 RA and 1 OA synovial tissue samples were collected and cryopreserved before being thawed, disaggregated, and FACS-sorted as described above.

### Unimodal scATAC-seq experimental protocol

Unimodal scATAC-seq experiments were performed by the BWH Center for Cellular Profiling. Each sample was processed separately in the cell capture step. Nuclei were isolated using an adaptation of the manufacturer’s protocol (10X Genomics). Approximately ten thousand nuclei were incubated with Tn5 Transposase. The transposed nuclei were then loaded on a Chromium Next GEM Chip H and partitioned into Gel Beads in-emulsion (GEMs), followed by GEM incubation and library generation. The ATAC libraries were sequenced to an average of 30,000 reads per cell with recommended number of cycles according to the manufacturer’s protocol (Single Cell ATAC V1.1, 10X Genomics) using Illumina Novaseq. Samples were initially processed using 10x Genomics Cell Ranger ATAC 1.1.0, which includes barcode processing and read alignment.

### Multiome experimental protocol

Multiome experiments were performed by the BWH Center for Cellular Profiling. Each sample was processed separately in the cell capture step. Nuclei were isolated as above. Approximately ten thousand nuclei transposed nuclei were loaded on Chromium Next GEM Chip J followed by GEM generation. 10x Barcoded DNA from the transposed DNA (for ATAC) and 10x Barcoded, full-length cDNA from poly-adenylated mRNA (for Gene Expression) were produced during GEM incubation. The ATAC libraries and Gene Expression libraries were then generated separately. Both library types were sequenced to an average of 30,000 reads per cell on different flowcells with recommended sequencing cycles according to the manufacturer’s protocol (Chromium Next GEM Single Cell Multiome ATAC + Gene Expression, 10X Genomics) using Illumina Novaseq. Samples were initially processed using 10x Genomics Cell Ranger ARC 2.0.0, which includes barcode processing and read alignment, for both ATAC and GEX information.

### ATAC quality control

The unimodal scATAC and multimodal snATAC datasets were processed separately, but in the same manner unless otherwise stated. Reads were quality controlled from the Cell Ranger BAM files via a new cell-aware strategy that removes likely duplicate reads from PCR amplification bias within a cell while keeping reads originating from the same positions but from different cells. For unimodal scATAC-seq data, duplicate reads from the same cell were called based on read and mate start positions and CIGAR scores, but the multimodal snATAC-seq data only used start positions since Cell Ranger ARC did not provide a mate CIGAR score (MC:Z flag). Reads that were not properly mapped within a pair, had a MAPQ < 60, did not have a cell barcode, or were overlapping the ENCODE blacklisted regions^24^ of ‘sticky DNA’ were also removed. BAM read files were converted to fragment BED files using BEDOPS^113^ bam2bed while accounting for the 9-bp Tn5 binding site. We kept cells with more than 10,000 reads with at least 50% of those reads falling in peak neighborhoods (5x full peak size), at least 10% of reads in promoter regions, not more than 10% of reads calling in the mitochondrial chromosome, and not more than 10% of pre-deduplication reads falling in the ENCODE backlisted regions^24^. The genome annotation we used to define promoters was GENCODE v28 basic^26^ as was done for Cell Ranger ATAC read mapping; we defined promoter regions for the QC step as 2kb upstream of HAVANA protein coding transcripts that we subsequently merged to avoid double counting. The fragments from the post QC cells were quantified within the 200bp trimmed consensus peaks (see **ATAC peak calling**) via GenomicRanges::findOverlaps^114^ into a peaks x cells matrix. We then did an initial round of broad cell type clustering: binarize peaks x cells matrix, log(TFxIDF) normalization using Seurat::TF.IDF^115^, most variable peak feature selection using Symphony::vargenes_vst^94^, center/scale features to mean 0 and variance 1 across cells using base::scale, PCA dimensionality reduction to 20 PCs using irlba::prcomp_irlba, batch correction by sample using Harmony::HarmonyMatrix^27^, shared nearest neighbor creation using RANN::nn2 and Seurat::ComputeSNN^115^, and Louvain clustering using Seurat::RunModulatrityClustering^115^. For the unimodal scATAC-seq broad cell type processing, we chose peaks that had at least one fragment in at least five percent of cells, TFxIDF normalization using Seurat::TF.IDF^115^, and PCA to 20 PCs using irlba::prcomp_irlba with centering and scaling internally before continuing in the above steps. We visualized clusters using UMAP coordinates via umap::umap. We removed doublet clusters with multiple cell-type-specific marker peaks (see **Broad cell type clustering**), intermediate placement between broad cell type clusters in principal component space, high fragment counts, and high doublet scores determined per cell per donor by ArchR^32^. Note that this does not necessarily preclude doublets of the same cell type.

### ATAC peak calling

For consistent analysis, we used trimmed consensus peaks across all ATAC cells for all analyses unless otherwise stated. Peaks were called twice, before and after ATAC cell QC, to first provide general peak information to be used in cell QC step and then afterwards on the post QC cells to provide the final, refined peak set. Individual scATAC-seq donor BAM files were converted to MACS2^116^ BEDPE files using macs2 randsample, concatenated across donors, and then used to call peaks with macs2 callpeak --call-summits using a control file^117^ where ATAC-seq was done on free DNA to account for Tn5’s inherent cutting bias. The best sub-peak, as determined by signal value and q-value, was trimmed to 200 bp (summit ± 100bp) to localize the signal and avoid confounding any statistical analysis with peak length. Any overlapping peaks were removed iteratively, keeping the best sub-peak, to avoid double counting. We confirmed these scATAC-seq peaks were reasonable to use for the multiome snATAC-seq datasets, beyond just that the datasets were done on the same tissue type, as there was an average of 75% (n=12 datasets; range: 66%-83%) of the 200bp trimmed snATAC-seq donor-specific peaks overlapping the scATAC-seq consensus peaks; we used the 5x full consensus peak neighborhoods in the cell QC step for multiome datasets as an added safeguard. We also confirmed our peaks’ quality by seeing good overlap with ENCODE SCREEN v3 candidate cis-regulatory elements (cCREs)^25^ and the GENCODE v28^26^ promoter annotations via bedtools^118^ intersectBed (**Supplementary Fig. 1f**).

### RNA quality control

snRNA cells had to pass Cell Ranger ARC cell filtering and have at least 500 genes and less than 20% of mitochondrial reads. The Cell Ranger ARC genes x cells matrix was subsetted to only these cells passing cell QC. We did an initial round of broad cell type clustering: log normalization to 10,000 reads using Seurat::NormalizeData^115^, most variable gene feature selection using a variance stabilizing transformation (VST)^115^, center/scale features to mean 0 and variance 1 across cells using base::scale, PCA dimensionality reduction to 20 PCs using irlba::prcomp_irlba, batch correction by sample via Harmony::HarmonyMatrix^27^, shared nearest neighbor creation using RANN::nn2 and Seurat::ComputeSNN^115^, and Louvain clustering using Seurat::RunModulatrityClustering^115^. We visualized clusters using UMAP coordinates using umap::umap. We removed doublet clusters with multiple cell-type-specific genes (see **Broad cell type clustering**), intermediate placement between broad cell type clusters in principal component space, high UMI counts, and high doublet scores determined per cell per donor by Scrublet^119^. Note that this does not necessarily preclude doublets of the same cell type.

### Broad cell type clustering

The unimodal scATAC and multimodal snATAC datasets were processed separately, but in the same manner unless otherwise stated. For cells passing QC, we subsetted the feature x cells matrices and preformed broad cell type clustering within modalities as described above. Marker peaks/genes denoting cell types were used as follows: *CD3D* and *CD3E* in T cells; *NCAM1* and *NCR1* in NK cells; *MS4A1* and *TNFRSF17* in B/plasma cells; *CD163* and *C1QA* in myeloid cells; *PDPN* and *PDGFRB* in fibroblasts; and *VWF* and *ERG* in endothelial cells. Marker peaks were defined as peaks overlapping the promoters of marker genes; if there were multiple peaks overlapping a gene’s promoter or multiple isoforms of a gene, the peak that best tracked with the gene’s expression in the multiome cells was chosen. We also classified the multiome snRNA cells into the AMP-RA CITE-seq study^12^ broad cell types using Symphony^94^ (see **Symphony classification of transcriptional cell state**). The small minority of cells (2%) with discordant cell types defined in the snATAC, snRNA, and CITE-seq modalities for the multiome datasets were removed. Here, as in all analyses, we included OA samples to increase cell counts, but we did not make any OA versus RA comparisons due to low power.

### Fine-grain chromatin class clustering

To define chromatin classes within broad cell types, we made peaks x cells matrices for each broad cell type combining unimodal scATAC-seq and multimodal snATAC-seq cells. Since peaks were called on all scATAC-seq cells regardless of cell type, we first subset each peaks x broad cell type cells matrix by “peaks with minimal accessibility” (PMA). We defined minimal accessibility as peaks that had a fragment in at least 0.5% of cells, except for endothelial cells which we increased to a minimum of 50 cells. After subsetting the matrix by PMA peaks, we ran the same clustering pipeline detailed in the broad cell type clustering section with 10 PCs requested. For T, stromal, myeloid, and B/plasma cell types, we used Harmony^27^ for batch-correction by sample with all other default parameters. For endothelial cells, due to small cell counts, we batch-corrected on both sample and assay and updated Harmony’s sigma parameter to 0.2. We did another round of QC to exclude cells that clustered primarily due to relatively fewer total fragments per cell and fewer peaks with at least one 1 fragment per cell, and then re-clustered. We tried a number of clustering resolutions (see **Supplementary Fig. 10** for a subset) and chose the resolution at which we could define clusters biologically with known markers that tracked in both chromatin accessibility and gene expression spaces.

### T cell lineage analysis

We used a logistic model to investigate how promoter peaks align with the CD4 and CD8 lineage distinction (‘lineage’) across cells beyond their chromatin class identity (‘class’), sample’s donor (‘donor’), and overall fragment counts (‘nFragments’). The lineage variable was defined as the cell’s chromatin accessibility at the promoter peaks of: CD4+ CD8A-(+1), CD4+ CD8A+ or CD4-CD8A-(0), CD4-CD8A+ (-1); cell counts by lineage and class are in **Supplementary Table 2**. Genome-wide T cell promoter peaks were defined as those T cell PMA peaks that overlapped an ENCODE promoter-like cCRE^25^, whose proposed target gene was assessed via overlapping ENSEMBL^120^ hg38 release 92 transcript annotations. For each of these binarized promoter peaks (‘peak’), we calculated two logistic regressions using lme4::glmer^121^:

Full model: peak ∼ lineage + class + (1|donor) + scale(log10(nFragments))

Null model: peak ∼ class + (1|donor) + scale(log10(nFragments))

A lineage beta in the model is positive if the peak is associated to CD4 and negative if associated to CD8. We calculated significance as a likelihood ratio test (LRT) between the full and null models with multiple hypothesis test correction using FDR<0.20; significant results are shown in **Supplementary Table 3**. Furthermore, we defined a lineage score by cell via: 1) subsetting the normalized chromatin accessibility matrix by the lineage-significant peaks; 2) dividing CD4-associated peaks by the number of CD4-associated peaks to normalize; 3) dividing CD8A-associated peaks by the number of CD8A-associated peaks to normalize; 4) multiplying CD8A-associated peaks by -1 to differentiate lineage; 5) summing over peaks by cell to get a cell score. Thus, if a cell’s lineage score is positive, that cell is more associated to CD4 and CD8 if otherwise. We aggregated these cell scores by chromatin class in **Supplementary Fig. 2d**.

### Transcription Factor motif analysis

We used ArchR^32^ version 1.0.2 for our TF motif analysis. For each cell type’s final QC cells, we subsetted each donor’s fragments using awk^122^, bgzip^123^, and tabix^124^ before creating arrow files from them using createArrowFiles with all additional QC flags nullified. ArchR removed samples with two or fewer cells, so one sample with only two B/plasma cells was removed in that cell type. From the arrow files, we created an ArchR project via ArchRProject. We added our peak set into the project by addPeakSet and recreated a peaks by cells matrix via addPeakMatrix. We added our chromatin classes to the project’s cell metadata with addCellColData. Then, we added motif annotations to our peaks using addMotifAnnotations with the JASPAR2020 motif set version 2, a 4 bp motif search window width, and motif p-value of 5e-05. We added chromVAR background peaks via addBgdPeaks and then calculated chromVAR deviations using addDeviationsMatrix. Next, we found marker peaks for each chromatin class using getMarkerFeatures via a Wilcoxon test and accounting for TSS Enrichment and log10(nFragments). Within those marker peaks, we found motif enrichment via peakAnnoEnrichment with cutoffs FDR <= 0.1 and Log2FC >= 0.5. We modeled our heatmap of motif enrichment on plotEnrichHeatmap, but we added some filters. As in the default plotEnrichHeatmap method, we used the -log10(padj), where the p-value is calculated via a hypergeometric test, as the motif enrichment value. For each chromatin class sorted by maximum motif enrichment value, we chose the top motifs not already chosen that had at least an enrichment value of 5 for that class, had the maximal or within 95% of the maximal enrichment for that class, and whose corresponding TF had at least 0.05 mean-aggregated normalized gene expression for that class. For myeloid cells, the enrichment cutoff was set to 2 to show some motifs for M_A_-0. In endothelial cells, there were so few E_A_-3 cells that only 1 marker peak was called for that class, resulting in no useful motif information to be shown; we also added a SOX17 motif (JASPAR^98^ ID MA0078.1), a prominent arteriolar endothelial TF^85^, to the JASPAR2020 motif set for endothelial cells. For the chosen motifs, we plotted the percentage of the max enrichment value across classes with the max value in parentheses in the motif label as in plotEnrichHeatmap.

### Loci visualization

To visualize the ATAC read buildups by chromatin class or transcriptional cell state (class/state), we first subsetted the deduplicated BAM files for each donor by the cells in the specific state/class using an awk^122^ command looking for the samtools CB:Z (*i.e.*, cell barcode) flag; a BAM index file was made for each BAM file for region subsetting purposes later. Then for each class/state at each locus, we subsetted each donor’s BAM file for that region using samtools view, merged the BAM files across donors using samtools merge, converted the BAM files to bedgraph files using bedtools^118^ genomecov, and then divided the bedgraph counts by the total read count (by 1e7 reads) in that class/state to allow for comparison between classes/states. The bedgraph files were then imported to IGV^125^ and the data range for each class/state was set to the maximum value across classes/states. Tracks were colored by their class/state. We did not always show all classes/states for space reasons, but we picked representatives that were similar in the locus shown. Peaks (see **ATAC peak calling**), motifs (see **Transcription Factor motif analysis**), and SNPs (see **Genetic variant analysis**) were imported into IGV as BED files. We could not label all motifs found in these loci for space reasons, so we picked the enriched motif we were highlighting and a few other motifs enriched in the highlighted class. We also could not always show all the gene isoforms for all loci for space reasons, but we did always show a representative isoform for those that looked similar in the locus shown.

### Stromal DNA methylation analysis

We downloaded 1859 differentially methylated (DM) loci for RA versus OA synovial fibroblast cell lines from Nakano et al., 2013^47^. We converted the 1 bp DM regions from hg19 to hg38 reference genomes using liftOver^126^; 1 region did not map. Next, we overlapped these DM loci with our 200 bp stromal PMA peaks using intersectBed^118^ to get 152 DM loci, 67 associated to hypermethylation and 85 to hypomethylation. We defined a per-cell score as in the **T cell lineage analysis** section, but with positive scores corresponding to hypermethylation and negative scores to hypomethylation. We calculated a Wilcoxon Rank Sum Test p-value of DNA methylation cell scores between the 11,733 cells in S_A_-0 and the 12,574 cells not in S_A_-0 to get significance.

### Tissue and blood analysis

We downloaded a publicly available 10x Single Cell Multiome ATAC + Gene Expression dataset^90^ of healthy donor (female, age 25) PBMCs with granulocytes removed through cell sorting as part of our sister study^91^ (‘Public PBMC’ dataset). The PBMC cell labels were generated using the processing defined in that study. No further quality control was done on the fragment file downloaded from the 10x website (https://cf.10xgenomics.com/samples/cell-arc/2.0.0/pbmc_granulocyte_sorted_10k/pbmc_granulocyte_sorted_10k_atac_fragments.tsv.gz). For each cell type (B, T, and myeloid), we subset the fragment file by that cell type’s cells and then overlapped them with our peaks to get a peaks x cells matrix as done in **ATAC quality control**. We concatenated this matrix to our RA tissue’s peaks x cells matrix for each corresponding cell type and then re-clustered using the same PMA and variable peaks chosen for tissue and harmonizing by sample. We chose the resolution that best mirrored the RA tissue chromatin classes. The odds ratio for each individual biological source’s cell label and the combined tissue and blood cluster label was calculated as in **Class/state odds ratio**.

### Symphony classification of transcriptional cell state

To determine the RA transcriptional cell states within our multimodal data, we used Symphony^94^ to map the multimodal snRNA profiles into the AMP-RA reference synovial tissue transcriptional cell states^12^. We used a Symphony reference object from that study for each broad cell type we tested (T cell, stromal, myeloid, B/plasma, and endothelial); the lymphocyte states were defined using both gene and surface protein expression while the others were defined using gene expression only. For each cell type, we mapped each multimodal snRNA gene x cells matrix into the appropriate Symphony reference object using the mapQuery function, accounting for donor as a batch variable. Using the knnPredict function with k=5, each multiome cell was classified into a reference transcriptional cell state by the most common annotation of its five nearest AMP-RA reference neighbors in the harmonized embedding. We considered it a high confidence mapping if at least 3 out of the 5 nearest reference neighbors were the same cell state, though the number of cell states will affect this as more cell states means more boundary regions between cell states.

### Class/state odds ratio

For each combination of chromatin class and transcriptional cell state within a cell type, we constructed a 2×2 contingency table of the number of cells belonging or not to the class and/or state. For cell states that had more than 10 classified cells, we then calculated the odds ratio (OR) and p-value via stats::fisher.test. We did multiple hypothesis test correction via stats::p.adjust using FDR<0.05. We displayed the natural log of the OR via base::log, and if the value was infinite, we capped it at 1 plus the ceiling of the non-infinite max absolute value of logged OR for display purposes; negative infinity was the negative capped number. All the ORs and p-values for all class/state combinations from **Fig. 7** and **Supplementary Fig. 8g-h** are in **Supplementary Table 4**.

### Linear discriminant analysis

We used linear discriminant analysis (LDA) to determine how well knowing the ATAC harmonized principal component (hPC) information helped predict the mRNA fine-grain cell states for each pairwise combination of states. We specifically use pairwise combinations instead of 1 versus all comparisons to assess the chromatin accessibility data’s ability to give rise to one or multiple transcriptional cell states. For each pair of transcriptional cell states within a broad cell type, we subset all data structures by those cells and remade the cell state vector into a 1-hot encoding. If either cell state of the pair has less than 50 cells, we excluded it from further analysis. We used the ten ATAC hPCs from the fine-grain chromatin class clustering (see **Fine-grain chromatin class clustering**). Covariates of donor (1-hot encoded for 12 donors) and scaled logged number of fragments (nFragments) were used since both can affect cell type identity. We trained an LDA model using MASS::lda on 75% of cells across the pair of states, verifying that the training and testing sets had cells from both states:

LDA model: cell state ∼ ATAC hPCs + donors + scale(log10(nFragments))

We tested the model using stats::predict for the 25% of held-out data and quantified the discriminative value of the model using an area under the curve AUC metric from ROCR^127^ library functions ROCR::prediction and ROCR::performance. Pairs of distinct clusters were only calculated once; the square matrices of results have the triangles mirrored. If the cell states were the same and a model was not run (identity line) or the model between pairs of clusters had a constant variable due to donors with too few cells (non-identity line), the box is greyed out.

### Symphony classification of chromatin class

To utilize the richer clinical information in the more abundant AMP-RA reference datasets, we classified each AMP-RA reference cell into a chromatin class. We used the same shared transcriptional spaces by cell type defined in **Symphony classification of transcriptional cell state**, but we reversed the reference and query objects in the knnPredict function, such that the multiome cells were in the ‘reference’ and the AMP-RA reference cells were in the ‘query’. We used the most common annotation of the 5 nearest multiome neighbors to classify the chromatin class in the AMP-RA reference cells. We averaged the 5 nearest multiome neighbors’ UMAP dimensions to visualize the classified chromatin classes in the AMP-RA reference cells on the ATAC-defined UMAPs.

### scATAC-seq and CITE-seq shared donor analysis

There were different samples that came from the same donors in the unimodal scATAC-seq and AMP-RA reference CITE-seq datasets. We expected similar, but not the same, chromatin class proportions for samples coming from the same donor’s tissue but put through different experimental protocols and class assignment methods. First, we filtered out any donors that did not have at least 200 scATAC or CITE cells in all cell types except endothelial in which we lowered the threshold to 100 cells. We then calculated the proportion of each sample’s cells coming from each chromatin class for each technology and plotted the CITE proportion by scATAC proportion for each donor, faceted by chromatin class in **Fig. 8a** and **Supplementary Fig. 12a**. We calculated the Pearson correlation and p-value for each chromatin class by stats::cor.test.

### Co-varying neighborhood analysis (CNA)

We used the significant CNA^95^ correlations between AMP-RA reference cell neighborhoods and sample-level covariates from our AMP-RA reference study^12^. We re-plotted the AMP-RA reference cell CNA correlations on the ATAC-defined UMAPs and re-aggregated them by classified chromatin class calculated in **Symphony classification of chromatin class**.

### Genetic variant analysis

We used the set of RA-associated non-coding SNP locations and statistically fine-mapped post-inclusion probabilities (PIPs) from our previously published RA multi-ancestry genome-wide association meta-analysis study^104^. We subsetted the SNPs by PIP>0.1 and overlapped their locations with our peaks using intersectBed^118^. For the overlapping peaks, we plotted their normalized chromatin accessibility mean-aggregated by chromatin class and scaled in **Fig. 8d** with more description in **Supplementary Table 5**. To determine broad cell type specificity of a peak’s accessibility, we calculated a Wilcoxon Rank Sum Test 1-sided “greater” p value between the normalized, aggregated, scaled peak accessibility in the broad cell type’s classes versus those classes in the other broad cell types. Classes were considered accessible for that peak if the scaled mean normalized peak accessibility over 24 classes and 11 peaks, z, > 1. We plotted example loci in **Fig. 8e** and **Supplementary Fig. 13** as described in **Loci visualization**; we excluded some chromatin classes for space, but we kept the most accessible chromatin classes and at least one chromatin class from each cell type at each locus. The TF motif logos in **Fig. 8e** and **Supplementary Fig. 13** were downloaded from JASPAR motif database^98^ for accession IDs MA0517.1 (STAT1::STAT2), MA0039.4 (KLF4), and MA1483.1 (ELF2); they were not to scale, but the motif position the SNP disrupts is aligned to the SNP. We further aggregated ATAC reads by transcriptional cell state for visualization purposes in **Supplementary Fig. 14**.

## Data Availability

Raw and processed data will be available on public repositories upon acceptance.

## Code Availability

The code used to generate the results presented herein can be found on GitHub (https://github.com/immunogenomics/RA_ATAC_multiome/).

## Acknowledgements

This work was supported by the Accelerating Medicines Partnership (AMP) in Rheumatoid Arthritis and Lupus Network. AMP is a public-private partnership (AbbVie Inc., Arthritis Foundation, Bristol-Myers Squibb Company, Foundation for the National Institutes of Health, GlaxoSmithKline, Janssen Research and Development, LLC, Lupus Foundation of America, Lupus Research Alliance, Merck Sharp & Dohme Corp., National Institute of Allergy and Infectious Diseases, National Institute of Arthritis and Musculoskeletal and Skin Diseases, Pfizer Inc., Rheumatology Research Foundation, Sanofi and Takeda Pharmaceuticals International, Inc.) created to develop new ways of identifying and validating promising biological targets for diagnostics and drug development. Funding was provided through grants from the National Institutes of Health (UH2-AR067676, UH2-AR067677, UH2-AR067679, UH2-AR067681, UH2AR067685, UH2-AR067688, UH2-AR067689, UH2-AR067690, UH2-AR067691, UH2AR067694, and UM2-AR067678). Accelerating Medicines Partnership and AMP are registered service marks of the U.S. Department of Health and Human Services. This work is supported in part by funding from the National Institutes of Health (1UH2AR067677-01, U01HG009379, UC2AR081023). We also acknowledge support by NIH NHGRI T32HG002295 and NIAMS T32AR007530 (to K. Weinand and A.N.); and NIH NIAMS AR078769 (to D.A.R.). S.S. was in part supported by the Uehara Memorial Foundation and the Osamu Hayaishi Memorial Scholarship. UK Birmingham is supported by the Versus Arthritis Research Into Inflammatory Arthritis Centre Versus Arthritis (Versus Arthritis grant 22072) and the EU Innovative Medicines Initiative RT CURE. We wish to thank Tiffany Amariuta, Kaitlyn A. Lagattuta, Anika Gupta, and Angela Zou for helpful discussion.

## Author Contributions

K. Weinand, S.S., and S.R. conceptualized the study. K. Weinand conducted all computational analyses. S.S., A.N., F.Z., and S.R. provided input on statistical analyses and study design. S.S., A.N., A.H.J., D.A.R., M.B.B., K. Wei, and S.R. provided input on cellular analysis and interpretation. S.S. and S.R. supervised the study. AMP RA/SLE Consortium recruited patients and obtained synovial biopsies for unimodal scATAC-seq. L.T.D. and K. Wei recruited patients for multimodal samples. K. Wei, A.H.J, G.F.M.W., A.N., and M.B.B. designed and implemented the tissue disaggregation, cell sorting, and single cell sequencing pipeline. A.H.J., K. Wei, and G.F.M.W supervised and executed the tissue disaggregation pipeline for unimodal scATAC-seq samples. K. Wei, G.F.M.W, and Z.Z. supervised and executed the tissue disaggregation pipeline for multimodal samples. K. Weinand, S.S., and S.R. wrote the initial manuscript. All authors contributed to editing the final manuscript.

## Competing Interests

S.R. is a founder for Mestag Therapeutics, a scientific advisor for Janssen and Pfizer, and a consultant for Gilead. D.A.R. reports personal fees from Pfizer, Janssen, Merck, GlaxoSmithKline, AstraZeneca, Scipher Medicine, HiFiBio, and Bristol-Myers Squibb, and grant support from Merck, Janssen, and Bristol-Myers Squibb outside the submitted work. D.A.R. is a co-inventor on the patent for Tph cells as a biomarker of autoimmunity.

