## Supplementary Figures for "The Chromatin Landscape of Pathogenic Transcriptional Cell States in Rheumatoid Arthritis"

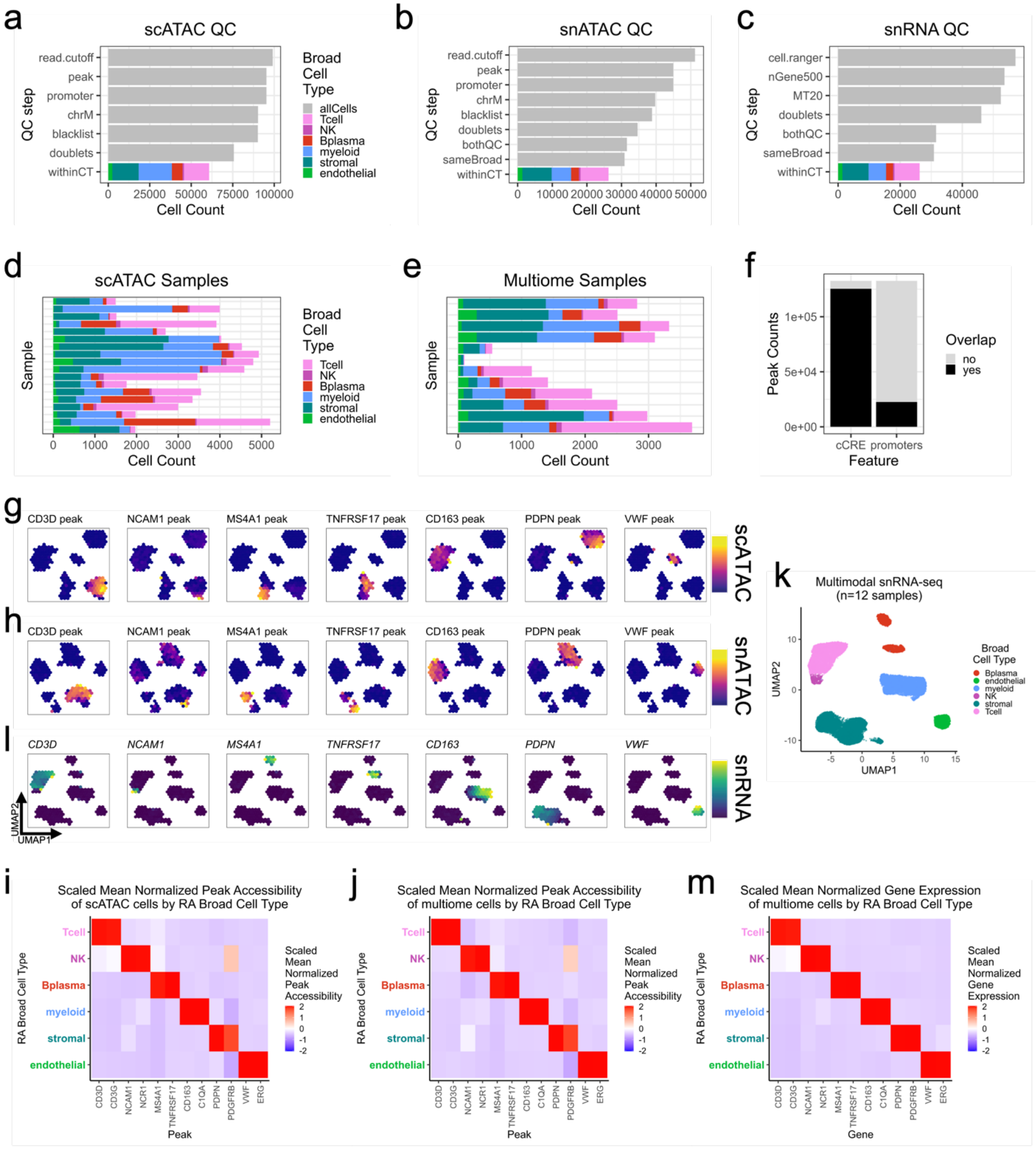


**Supplementary Fig. 1.** Quality Control Measures and Broad Cell Type Markers.
**a.-c.** Quality control steps ending in the final cell counts for all broad cell types in (**a.**) unimodal scATAC, (**b.**) multimodal snATAC, and (**c.**) multimodal snRNA datasets.
**d.-e.** Broad cell type cell counts per sample for (**d.**) unimodal and (**e.**) multimodal datasets.
**f.** Number of trimmed open chromatin peaks overlapping ENCODE cCREs[25] and promoters[26].
**g.-h**. Binned normalized marker peak accessibility visualized across broad cell types on the corresponding UMAPs for (**g.**) unimodal scATAC and (**h.**) multimodal snATAC datasets.
**i.-j.** Scaled mean normalized marker peak accessibility across broad cell types for (**i.**) unimodal scATAC and (**j.**) multimodal snATAC datasets.

**k.** Broad cell type identification for multiome snRNA cells.
**l.** Binned normalized marker gene expression visualized across broad cell types on the corresponding UMAP for multimodal snRNA datasets.
**m.** Scaled mean normalized marker gene expression across broad cell types for multimodal snRNA datasets.

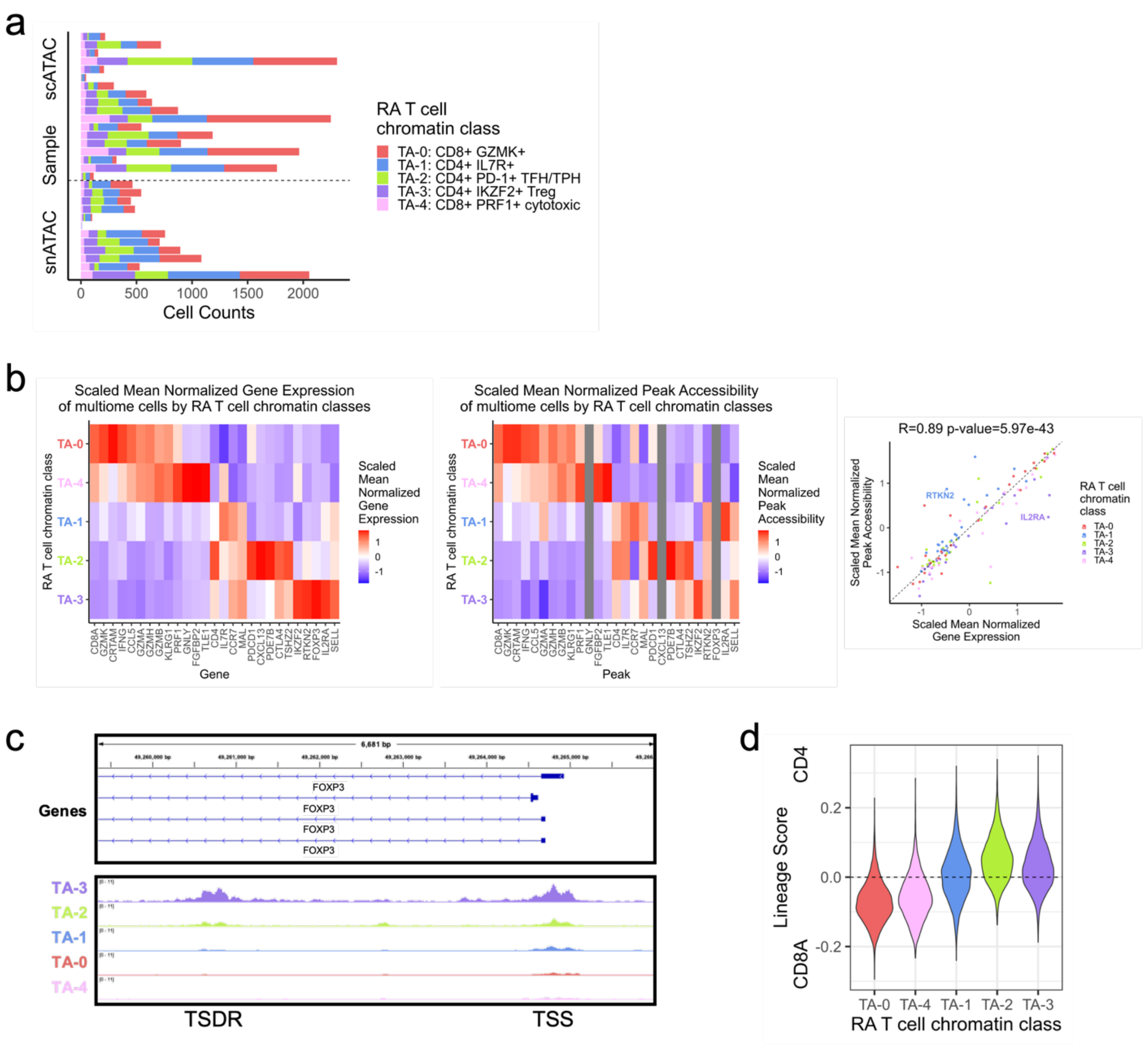


**Supplementary Fig. 2.** Further analyses of RA T cell chromatin classes.
**a.** T cell chromatin class cell counts per sample for unimodal and multimodal datasets.
**b.** Scaled mean normalized marker gene expression (**left**) and peak accessibility (**middle**) across T cell chromatin classes in multimodal datasets and the Pearson correlation (R) and p-value between them (**right**). Genes without an overlapping promoter peak are colored in grey in the middle panel. On the right, correlation calculated between 120 gene/class combinations.
**c.** FOXP3 locus (chrX:49,259,337-49,266,016) with selected isoforms and chromatin accessibility reads aggregated by chromatin class and scaled by read counts per class (**Methods**). Transcriptional start site (TSS) and Treg-specific demethylated region[29] (TSDR) highlighted as areas of selective open chromatin in $T_A$-3: CD4+ IKZF2 Tregs.
**d.** Lineage score by cell based on normalized peak accessibility for lineage-associated promoter peaks (**Methods**), segregated by T cell chromatin classes.

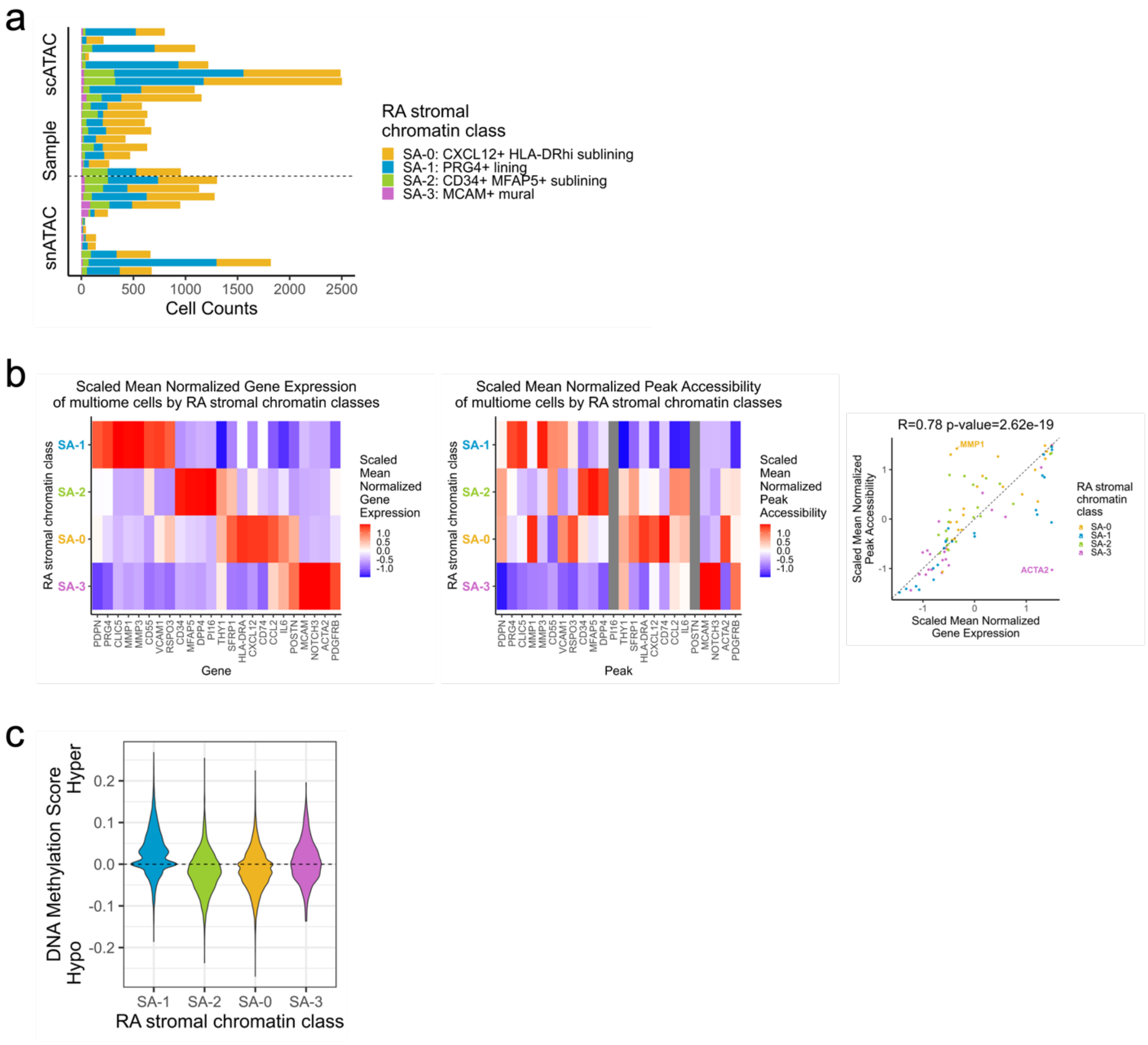


**Supplementary Fig. 3.** Further analyses of RA stromal chromatin classes.
**a.** Stromal chromatin class cell counts per sample for unimodal and multimodal datasets.
**b.** Scaled mean normalized marker gene expression (**left**) and peak accessibility (**middle**) across stromal chromatin classes in multimodal datasets and the Pearson correlation (R) and p-value between them (**right**). Genes without an overlapping promoter peak are colored in grey in the middle panel. On the right, correlation calculated between 88 gene/class combinations.
**c.** DNA methylation score by cell based on normalized peak accessibility for differential DNA methylation-associated promoter peaks (**Methods**), segregated by stromal chromatin classes.

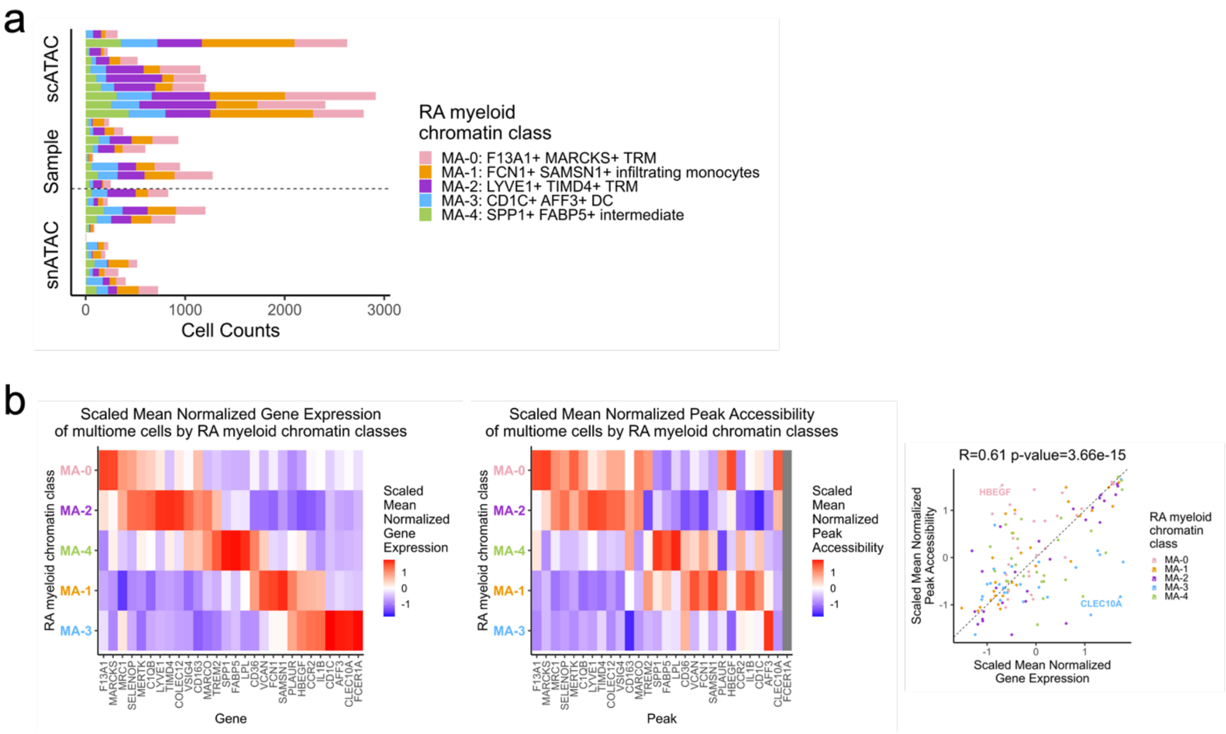


**Supplementary Fig. 4.** Further analyses of RA myeloid chromatin classes.
**a.** Myeloid chromatin class cell counts per sample for unimodal and multimodal datasets.
**b.** Scaled mean normalized marker gene expression (**left**) and peak accessibility (**middle**) across myeloid chromatin classes in multimodal datasets and the Pearson correlation (R) and p-value between them (**right**). Genes without an overlapping promoter peak are colored in grey in the middle panel. On the right, correlation calculated between 135 gene/class combinations.

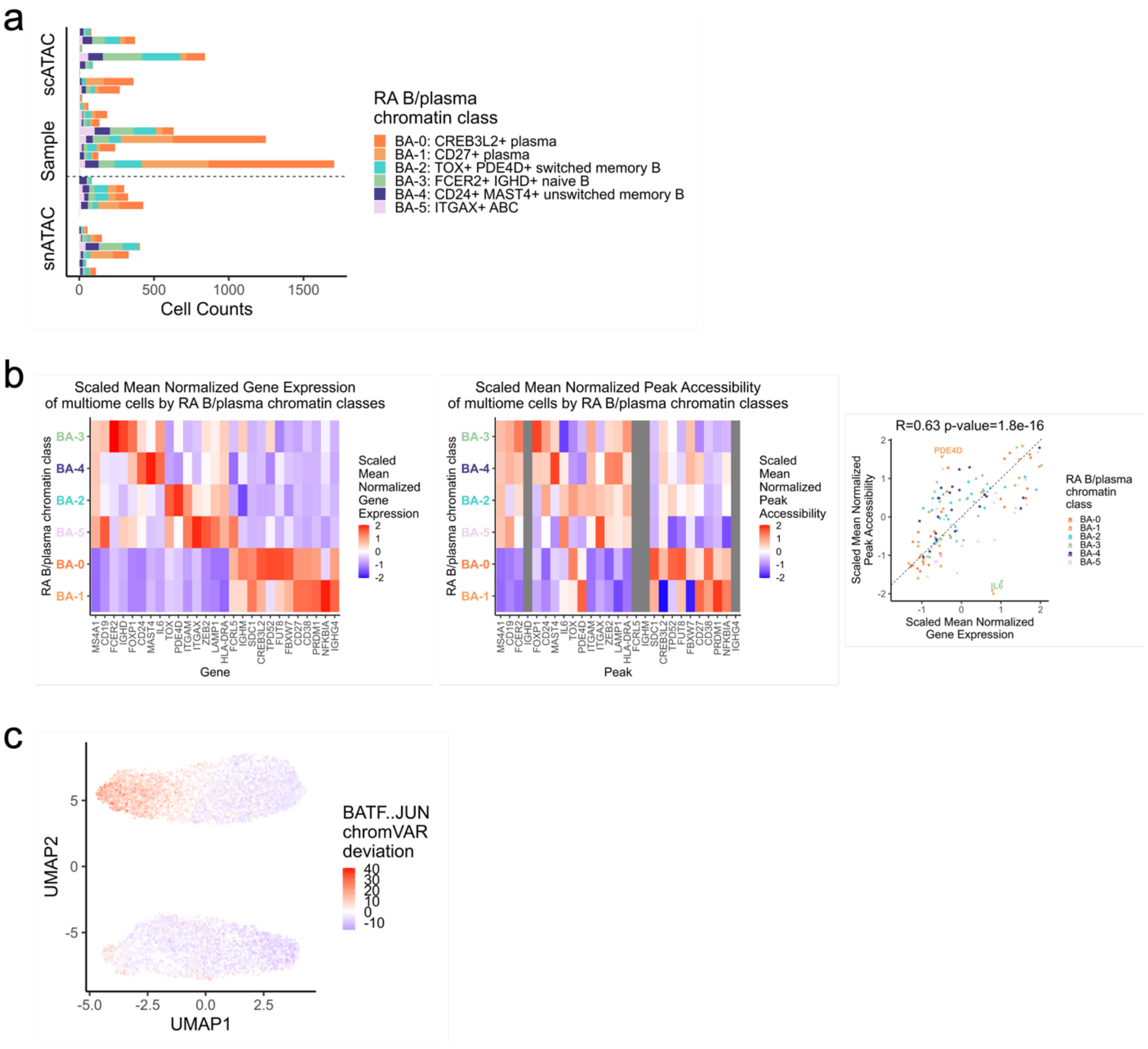


**Supplementary Fig. 5.** Further analyses of RA B/plasma chromatin classes.
**a.** B/plasma chromatin class cell counts per sample for unimodal and multimodal datasets.
**b.** Scaled mean normalized marker gene expression (**left**) and peak accessibility (**middle**) across B/plasma chromatin classes in multimodal datasets and the Pearson correlation (R) and p-value between them (**right**). Genes without an overlapping promoter peak are colored in grey in the middle panel. On the right, correlation calculated between 138 gene/class combinations.
**c.** UMAP colored by chromVAR[31] deviations for the BATF..JUN motif.

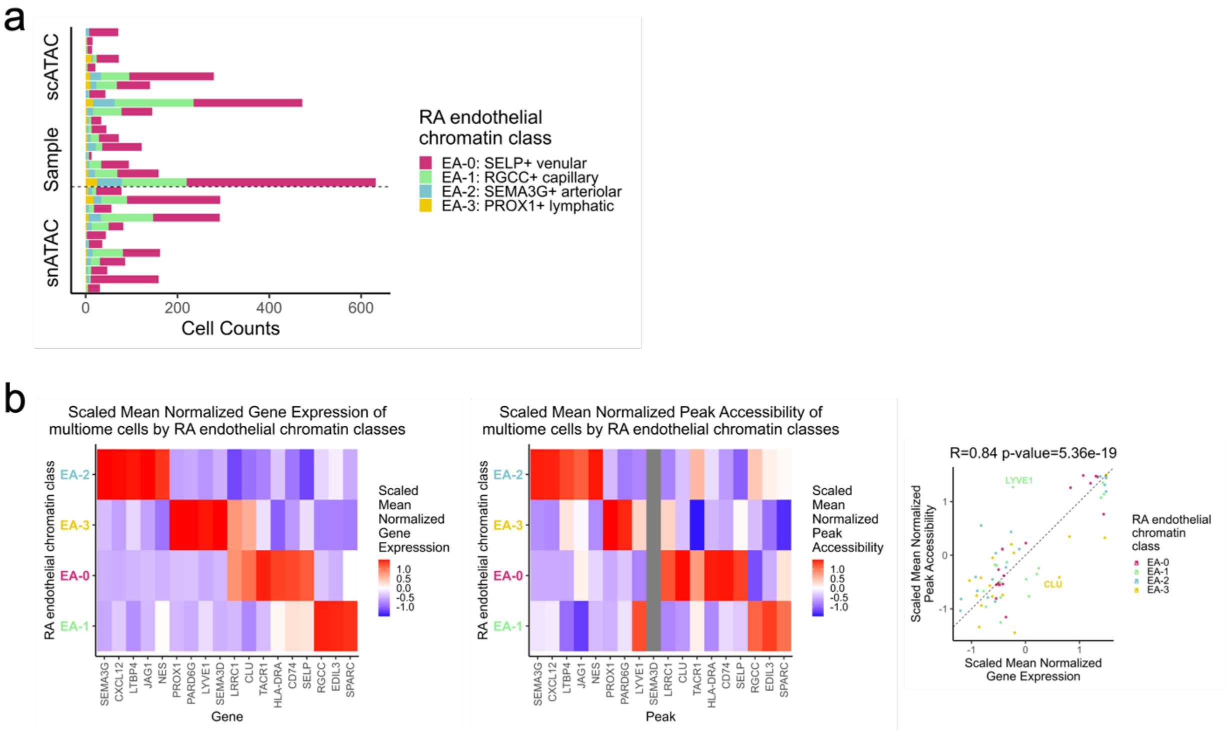


**Supplementary Fig. 6.** Further analyses of RA endothelial chromatin classes.
**a.** Endothelial chromatin class cell counts per sample for unimodal and multimodal datasets.
**b.** Scaled mean normalized marker gene expression (**left**) and peak accessibility (**middle**) across endothelial chromatin classes in multimodal datasets and the Pearson correlation (R) and p-value between them (**right**). Genes without an overlapping promoter peak are colored in grey in the middle panel. On the right, correlation calculated between 68 gene/class combinations.

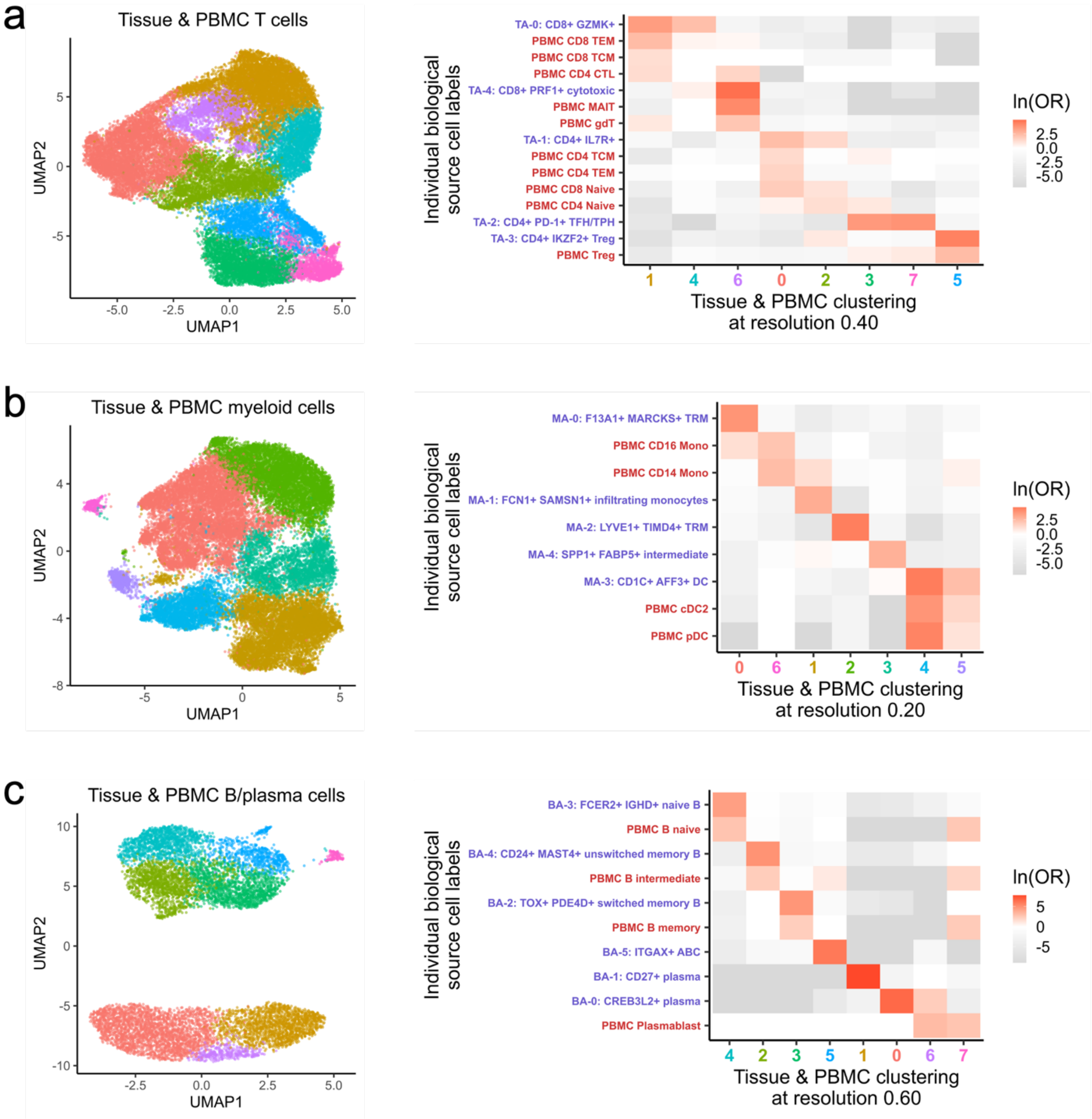


**Supplementary Fig. 7.** Tissue and PBMC analysis.
Clustering RA unimodal scATAC, RA multimodal snATAC, and healthy multimodal PBMC snATAC together visualized on UMAP (**left**) and natural log of Odds Ratio between these clusters and RA tissue/PBMC labels (**right**) for (**a.**) T, (**b.**) myeloid, and (**c.**) B/plasma cells. Non-significant (FDR>0.05) OR values are white. On the right, RA tissue chromatin classes are colored in purple and PBMC labels with more than 10 cells are colored in red; colors of x-axis labels correspond to the colors in the UMAPs on the left.

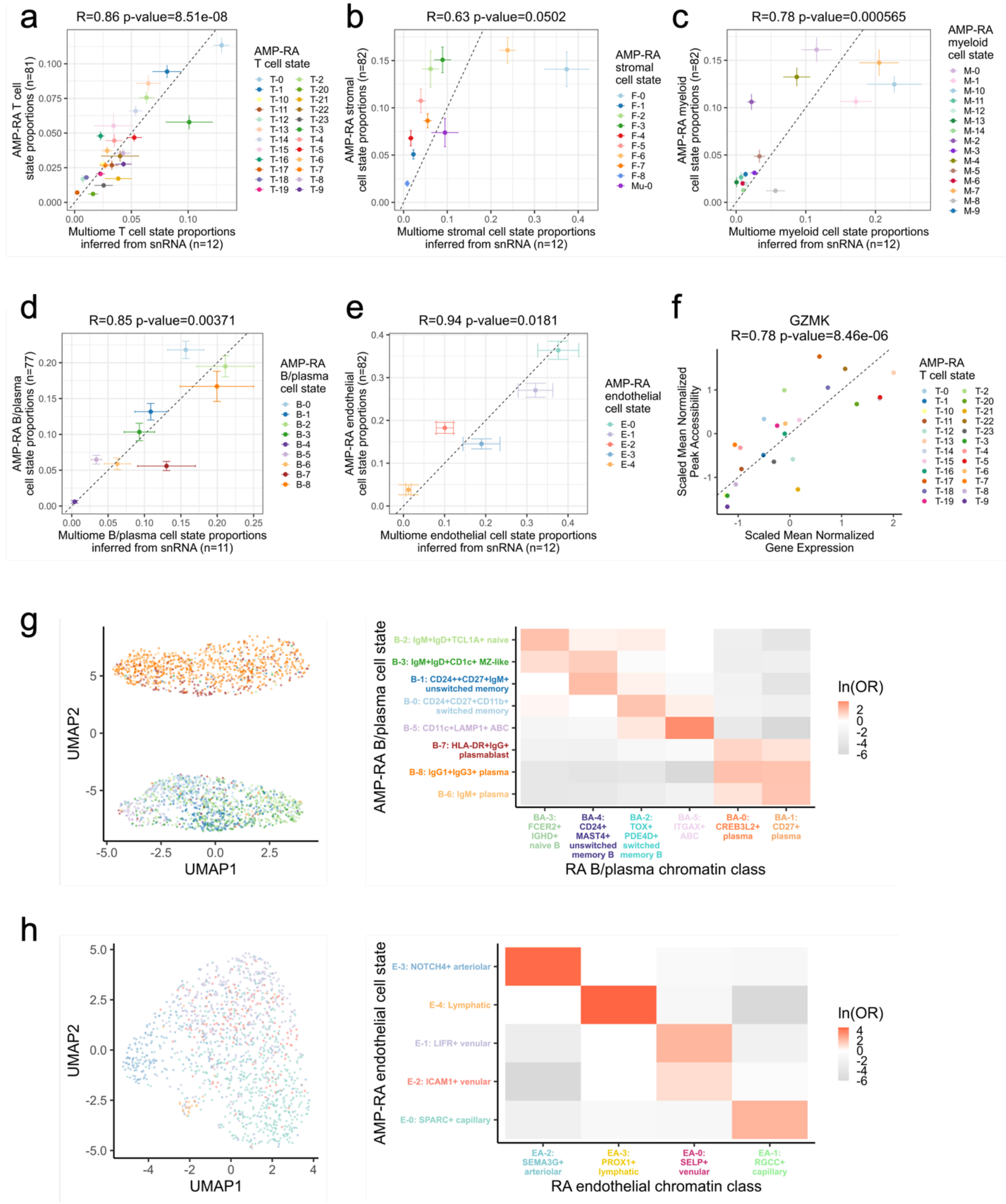


**Supplementary Fig. 8.** Mapping to transcriptional cell states and chromatin class superstate model.

**a.-e.** AMP-RA reference transcriptional cell state proportions across CITE datasets (y-axis) and multiome snATAC datasets (x-axis) for (**a.**) T (n=24), (**b.**) stromal (n=10), (**c.**) myeloid (n=15), (**d.**) B/plasma (n=9), and (**e.**) endothelial (n=5) cell states. Error bars are standard errors of the means. Pearson Correlation Coefficients (R) and p-values also noted.

**f.** Scaled mean normalized marker gene expression and peak accessibility for *GZMK* gene across AMP-RA reference transcriptional T cell states in multimodal datasets. Pearson correlation coefficient (R) and p-value calculated on the 24 T cell states.
**g.-h.** For (**g.**) B/plasma and (**h.**) endothelial cells, UMAP colored by classified AMP-RA reference transcriptional cell states for multiome cells (**left**) and natural log of Odds Ratio between chromatin classes and transcriptional cell states (**right**). Non-significant values (FDR>0.05) are white. In **g.**, B-4: AICDA+BCL6+ GC-like transcriptional cell state was excluded as fewer than 10 cells were classified into it.

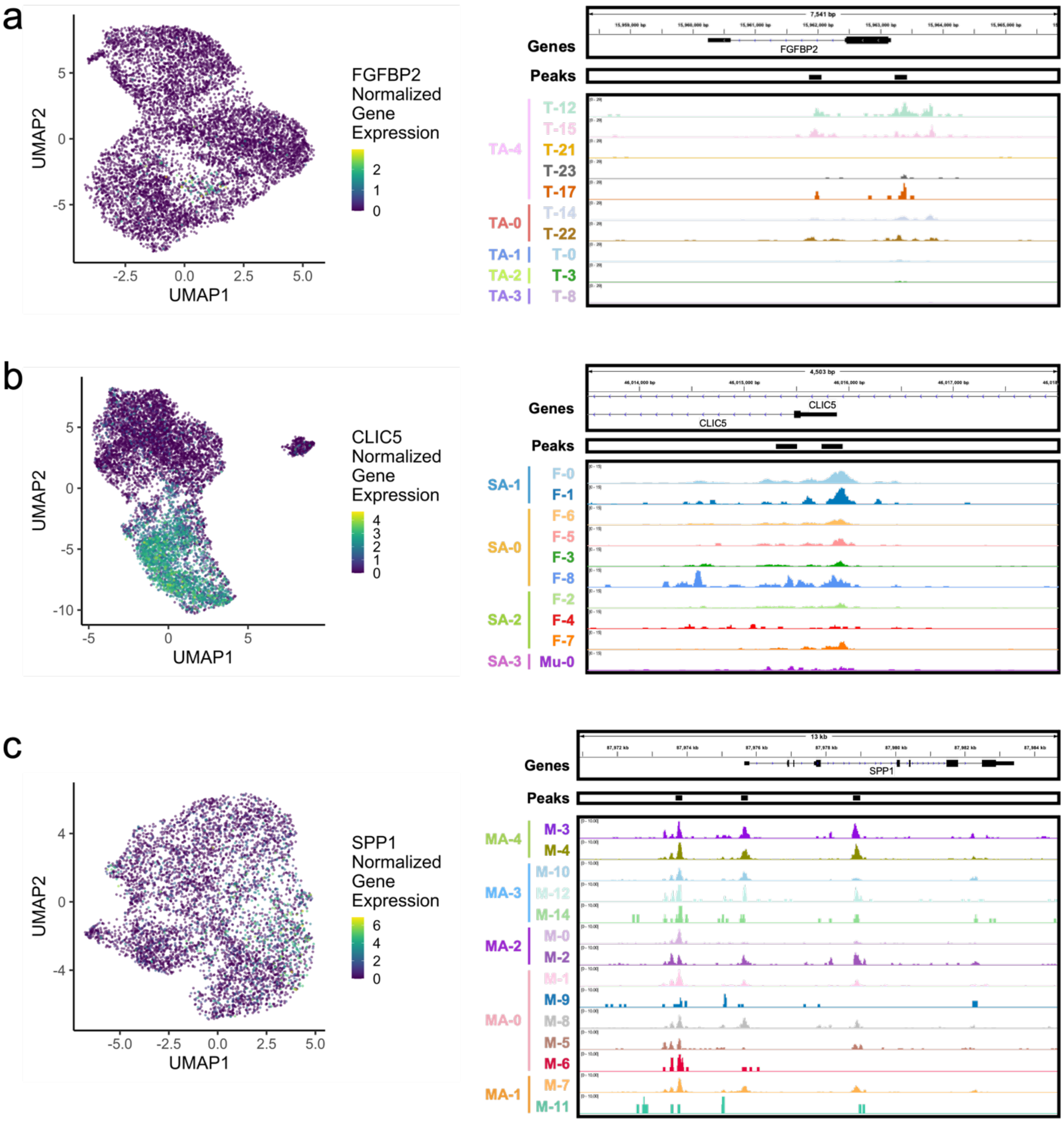


**Supplementary Fig. 9.** Open chromatin aggregated by transcriptional cell states.
UMAP colored by normalized gene expression of multiome cells (**left**) and gene loci with selected isoforms, open chromatin peaks, and chromatin accessibility reads aggregated by transcriptional cell state and scaled by cell counts per state (**Methods**) (**right**) for (**a.**) *FGFBP2* (chr4:15,958,313-15,965,852) in T cells, (**b.**) *CLIC5* (chr6:46,013,500-46,018,000) in stromal cells, and (**c.**) *SPP1* (chr4:87,970,884-87,984,690) in myeloid cells. The associated chromatin classes were also listed, though not all states therein were shown. In **c.**, M-13: pDC was excluded in myeloid cells since less than 10 multiome cells were classified as such.

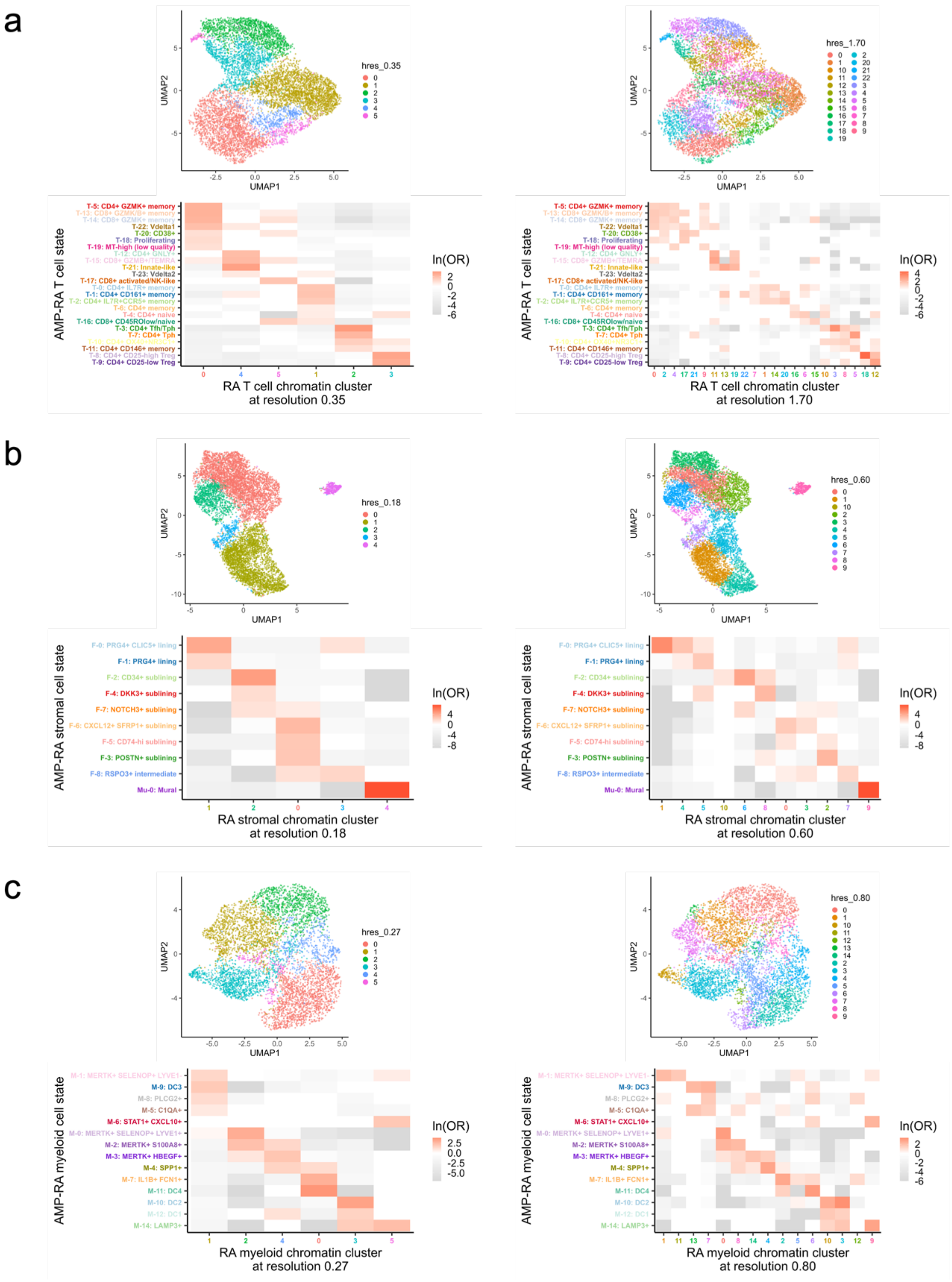

**Supplementary Fig. 10.** Additional ATAC clustering resolutions.
UMAP colored by ATAC clustering resolution (**top**) and natural log of Odds Ratio between ATAC clusters and transcriptional cell states (**bottom**) at clustering resolutions that add 1 additional cluster (**left**) and roughly the same number of clusters as states (**right**) for (**a.**) T, (**b.**) stromal, and (**c.**) myeloid cell types.

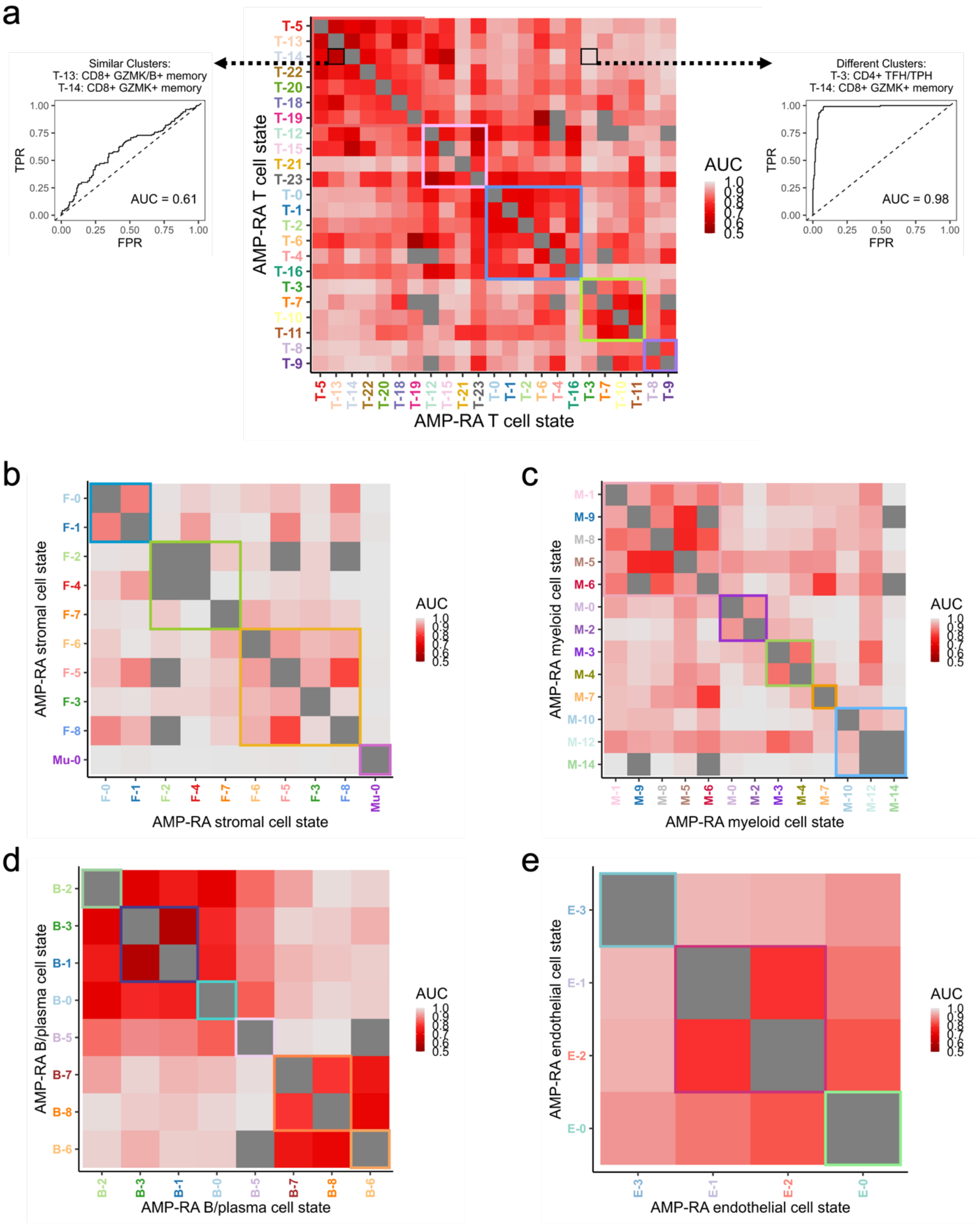


**Supplementary Fig. 11.** Linear Discriminant Analysis (LDA) model predicted transcriptional cell state from continuous ATAC principal components (PCs).
**a.** Linear Discriminant Analysis (LDA) model predicted the transcriptional cell state for each pair of T cell states from the ATAC principal components (PCs) (**Methods**). Area under the curve

(AUC) values shown for an example of similar states (**left**) and different states (**right**) with all pairs of states with more than 50 cells shown in the **middle**. Upper and lower triangles are mirrored; the LDA analysis was run only once per pair of states. The grey non-diagonal values depict pairs of states that had a constant variable in the model due to donors with too few cells. The superimposed colored boxes reflect chromatin class superstates defined in **Fig. 7** and **Supplementary Fig. 8g-h**.
**b.-e.** Middle panel of (**a.**) for (**b.**) stromal, (**c.**) myeloid, (**d.**) B/plasma, and (**e.**) endothelial cell types.

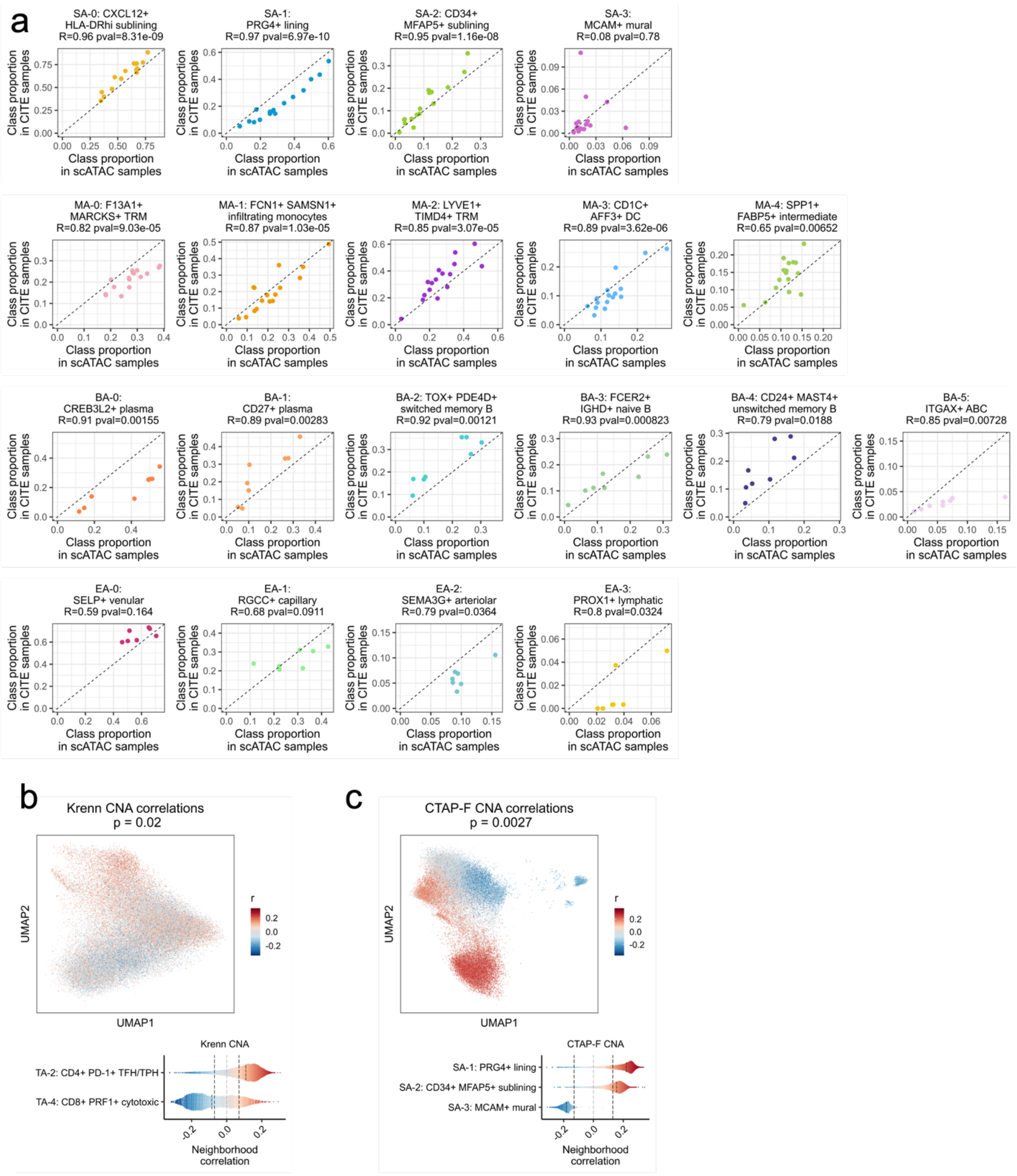


**Supplementary Fig. 12.** Additional RA CNA correlations.

**a.** For each donor shared between the unimodal ATAC and AMP-RA reference studies with at least 200 cells for that cell type (or 100 cells for endothelial cells), the Pearson correlation between the relative proportions of chromatin classes defined in the unimodal ATAC datasets (**x-axis**) and classified into in the CITE datasets through the multiome cells (**y-axis**). Pearson Correlation Coefficients (R) and p-values (pval) noted.

**b.** CNA correlations between T cell neighborhoods and Krenn inflammation score in AMP-RA

reference T cells visualized on UMAP (**top**) and aggregated by classified T cell chromatin classes (**bottom**). On the top, cells not passing the FDR threshold were colored grey. On the bottom, FDR thresholds shown in dotted black lines.
**c.** CNA correlations between stromal cell neighborhoods and CTAP-F in AMP-RA reference stromal cells visualized on UMAP (**top**) and aggregated by classified stromal chromatin classes (**bottom**). On the top, cells not passing the FDR threshold were colored grey. On the bottom, FDR thresholds shown in dotted black lines.

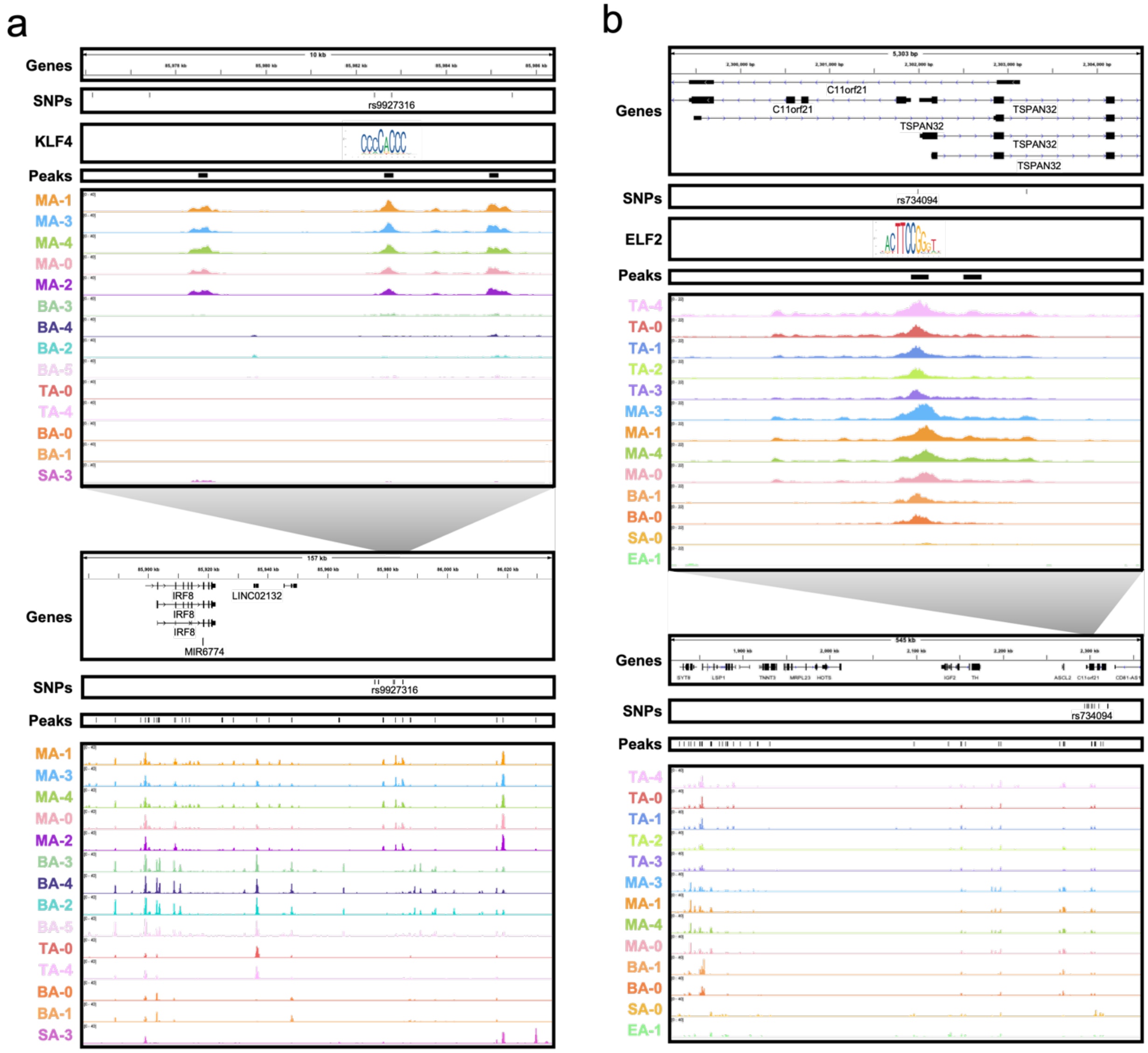


**Supplementary Fig. 13.** Chromatin accessibility of additional RA risk variants by chromatin classes.

**a.** rs9927316 locus, zoomed in (chr16:85,975,899-85,986,400) (**top**) and zoomed out (chr16:85,877,695-86,035,055) (**bottom**) with isoforms, SNPs, open chromatin peaks, and chromatin accessibility reads aggregated by chromatin class and scaled by read counts per class (**Methods**). KLF4 motif was downloaded from JASPAR[98] ID MA0039.4 and is not to scale, but it is aligned to the SNP-breaking motif position.

**b.** rs734094, zoomed in (chr11:2,299,199-2,304,500) (**top**) and zoomed out (chr11:1,815,000-2,360,000) (**bottom**) with isoforms, SNPs, open chromatin peaks, and chromatin accessibility reads aggregated by chromatin class and scaled by read counts per class (**Methods**). ELF2 motif was downloaded from JASPAR[98] ID MA1483.1 and is not to scale, but it is aligned to the SNP-breaking motif position.

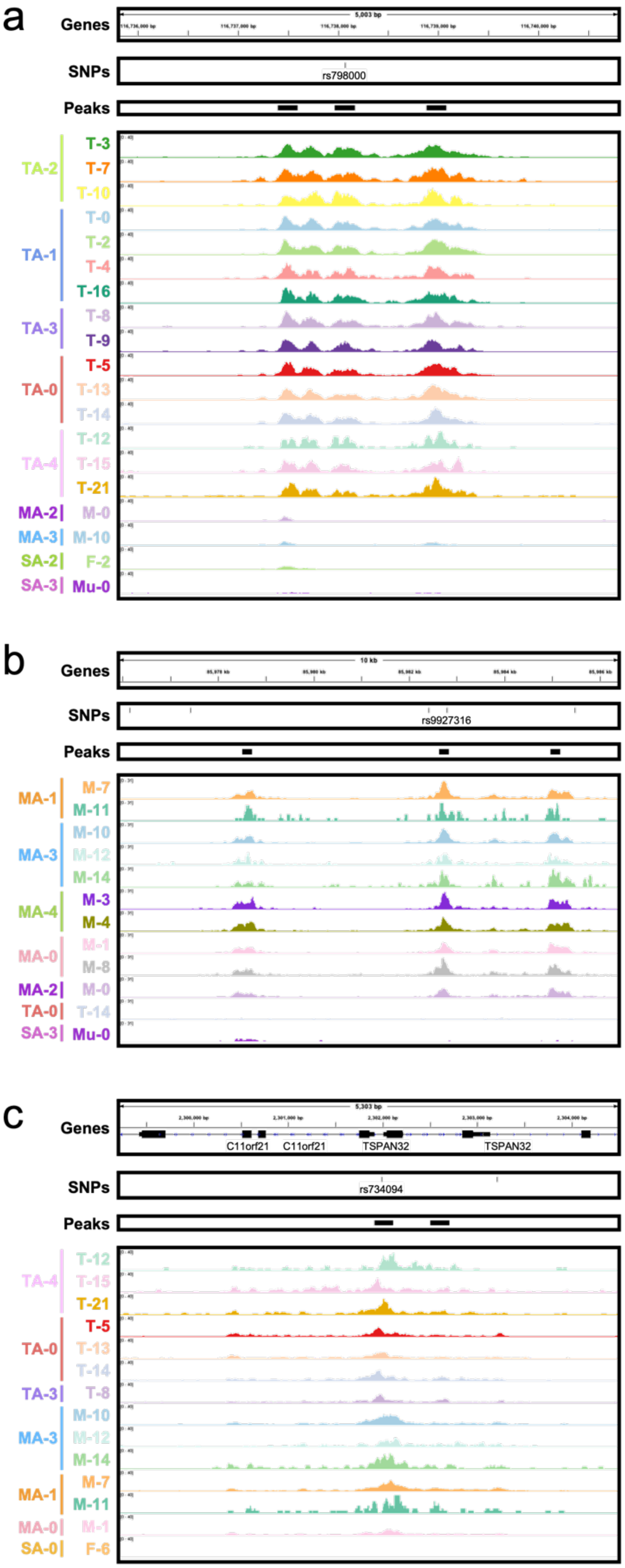


**Supplementary Fig. 14.** Chromatin accessibility of putative RA risk variants by transcriptional cell states.

Gene isoforms, SNPs, open chromatin peaks, and chromatin accessibility reads from multiome cells aggregated by transcriptional cell state and scaled by read counts per state (**Methods**) for (**a.**) rs798000 locus (chr1:116,735,799-116,740,800), (**b.**) rs9927316 locus (chr16:85,975,899-85,986,400), and (**c.**) rs734094 locus (chr11:2,299,199-2,304,500). The associated chromatin classes were also listed, though not all states therein were shown.
